# Complementation cloning identifies the essentials of mammalian Mastl kinase activation

**DOI:** 10.1101/2020.06.30.179580

**Authors:** Mehmet Erguven, Ezgi Karaca, M. Kasim Diril

## Abstract

Mastl is a mitotic kinase that is essential for error-free chromosome segregation. It is an atypical member of the AGC kinase family, possessing a unique non-conserved middle region (NCMR). The mechanism of its activation prior to mitosis has been extensively studied in Xenopus egg extracts. These studies found several residues (corresponding to T193 and T206 in the activation loop, and S861 in the C-terminal tail, i.e., C-tail of mouse Mastl) whose phosphorylations are crucial for enzymatic activation. To date, the significance of these phosphosites was not confirmed in live mammalian cells. Here, we utilize a complementation cloning approach to determine the essentials of mammalian Mastl kinase activity. We employed a tamoxifen-inducible conditional knockout system to delete the endogenous Mastl in mouse embryonic fibroblasts (MEF) and screened various mutants for their ability to complement its loss. MEFs, ectopically expressing different phosphorylation site mutants, were induced to undergo recombination-mediated knockout in their endogenous Mastl loci. S861A and S861D mutants were able to complement endogenous Mastl loss with proliferation rates comparable to WT. In parallel, we examined the available protein kinase structures having a phosphorylated C-tail. Among the published states, two distinct positionings of the C-tail phosphoresidue were observed. Energetic analysis of these states revealed that only one conformation highly contributes to the C-tail docking. Our in-depth sequence and structure analysis showed that Mastl pS861 does not belong to the conformational state, where the phosphoresidue contributes to the C-tail docking. The C-tail of Mastl is relatively short and it lacks the hydrophobic (HF) motif. In other AGC kinases, the C-tail phosphosite aids the anchoring of this motif over the N-lobe, leading to the final step of kinase activation. Together with the lack of HF motif in Mastl, our results suggest that phosphorylation of the C-tail turn motif phosphosite (S861) is auxiliary and is dispensable for mammalian Mastl kinase function. Furthermore, we demonstrated that complementation cloning is a powerful approach for screening the determinants of an essential protein’s functioning.

## 1. INTRODUCTION

### 1.1. Function of Mastl in mitosis

Mitosis is the most vibrant phase of cell cycle in which the genetic material of the parental cell is equally distributed to the daughter cells. It is characterized by dramatic and rapid alterations in the biochemical and the morphological state of the cell (Golloshi et al., 2017; Poon, 2007; Ramkumar & Baum, 2016). The initiation, progress, and termination of mitosis are strictly regulated by multiple layers of fail-safe regulatory mechanisms. These are post-translational modification, subcellular localization, and proteolytic degradation (Baxter et al., 2008). Post-translational regulation by phosphorylation is the major regulator of mitosis. Therefore, the initiation of mitosis is governed by mitotic protein kinases (H. Kim et al., 2016). Cdk1 is the primary kinase that governs mitosis (Diril et al., 2012; Santamaría et al., 2007). Key mitotic kinases work in a network, layered of positive feedback loops that construct a robust signaling cascade. This signaling cascade quickly activates the mitotic substrates. During the termination of the mitotic state at the beginning of anaphase, the mitotic signals are quenched by antagonistic protein phosphatases and the ubiquitin ligases. Actions of these antagonistic factors ultimately revert the cellular changes that occur during mitosis. The balance between the mitotic factors and the anti-mitotic factors is set on a cell cycle checkpoint named spindle assembly checkpoint (SAC). In general, while mitotic factors maintain SAC signaling, anti-mitotic factors suppress SAC in order to allow the cell to proceed to anaphase. Under normal circumstances, transition from metaphase to anaphase is blocked until all the kinetochores are correctly bound by spindle bilaterally (Ciliberto & Shah, 2009). SAC signaling preserves the genomic stability by delaying anaphase until each kinetochore-microtubule attachment is correct.

Cdk1 is present at the top of the mitotic signaling cascade. Once active, Cdk1 activates its activator Cdc25 (Yu et al., 2006) and inhibits its inhibitors Wee1 and Myt1 kinases. This positive feedback loop allows rapid accumulation of the active Cdk1 pool. Cdk1 and its downstream protein kinases activate their mitotic substrates, initiating a signaling cascade. Most of these phosphorylations on downstream proteins result in activation of factors that promote mitosis and inhibition of factors that promote mitotic exit. To preserve the mitotic state of the cell, mitotic phosphorylations have to be maintained throughout the phase. Therefore, a strong autoamplification loop has to be coupled with inhibition of the factors that promote mitotic exit (Gharbi-Ayachi et al., 2010). PP2A (protein phosphatase 2A) is the primary antagonist of Cdk1 during mitosis. This phosphatase is indirectly inhibited by Mastl (Microtubule-associated serine/threonine kinase like). Mastl is the human orthologue of Greatwall kinase (Gwl), first identified in Drosophila (Yu et al., 2004). Mastl kinase phosphorylates two highly homologous, small, thermostable proteins named Arpp19 and Ensa at their Serine 62 and Serine 67 residues, respectively. These two phosphoproteins specifically bind and inhibit B55∂ bound PP2A. Inhibition of PP2A-B55∂ results in rapid accumulation of phosphorylations on mitotic substrates. This inhibition also prevents removal of the inhibitory phosphorylations on Wee1 and Myt1, and the activating phosphorylation on Cdc25, further supporting the autoamplification loop and sustained MPF (maturation promoting factor) activity (M.-Y. Kim et al., 2012; Kishimoto, 2015; Mochida et al., 2010; Mochida & Hunt, 2012; Slupe et al., 2011).

Due to its crucial function in mitosis, Mastl is an essential protein kinase for cell division and proliferation. Loss-of-function studies have shown that, in the absence of Mastl, SAC signaling is weakened which results in premature onset of anaphase. Spindle-kinetochore attachment errors cannot be corrected, resulting in anaphase bridges and DNA breaks during chromosome segregation (Bisteau et al., 2020; Diril et al., 2016). The anaphase bridges prevent closure of the cleavage furrow, causing mitotic collapse at the end of mitosis (Alvarez-Fernandez et al., 2013).

### 1.2. Activation mechanism of AGC family protein kinases

Protein kinases are transferase class of enzymes that transfer the gamma phosphate of ATP to the serine, threonine, tyrosine, or histidine residues of the substrate proteins. Protein kinases are bilobal enzymes and the active site is buried in a cleft between its two lobes. The catalytically crucial segments or residues located at the catalytic cleft are the activation loop, DFG motif (within the activation loop), catalytic aspartate (within the DFG motif), glycine-rich loop, and a conserved lysine residue (at the N-lobe moiety of the ATP binding pocket).

The common kinase fold is constituted of a β-sheet-rich smaller N-lobe and an α-helix-rich larger C-lobe, connected through a loop segment. This loop encompasses the activation loop. DFG motif resides at the N-terminal of this connecting loop. The catalytic aspartate is located in the **<u>D</u>**FG (**<u>Asp</u>**-Phe-Gly) motif. Two divalent cations are accommodated into the ATP binding pocket in order to stabilize the negatively charged phosphate groups of ATP. The phosphate groups of ATP and the catalytic aspartate side chain together chelate cation cofactors, which are usually magnesium but can also be manganese (Diamant et al., 1995). The catalytic aspartate is responsible for the transfer of the ATP γ-phosphate to the target residue of the substrate protein or peptide. The glycine-rich loop’s backbone, chelated cofactors, and a conserved lysine together coordinate the ATP through charged or polar interactions with its triphosphate moiety. The purine moiety contributes to the coordination process by establishing specific hydrogen bonds with the hinge region backbone (Endicott et al., 2012) (Figure 1A).

**Figure 1.**
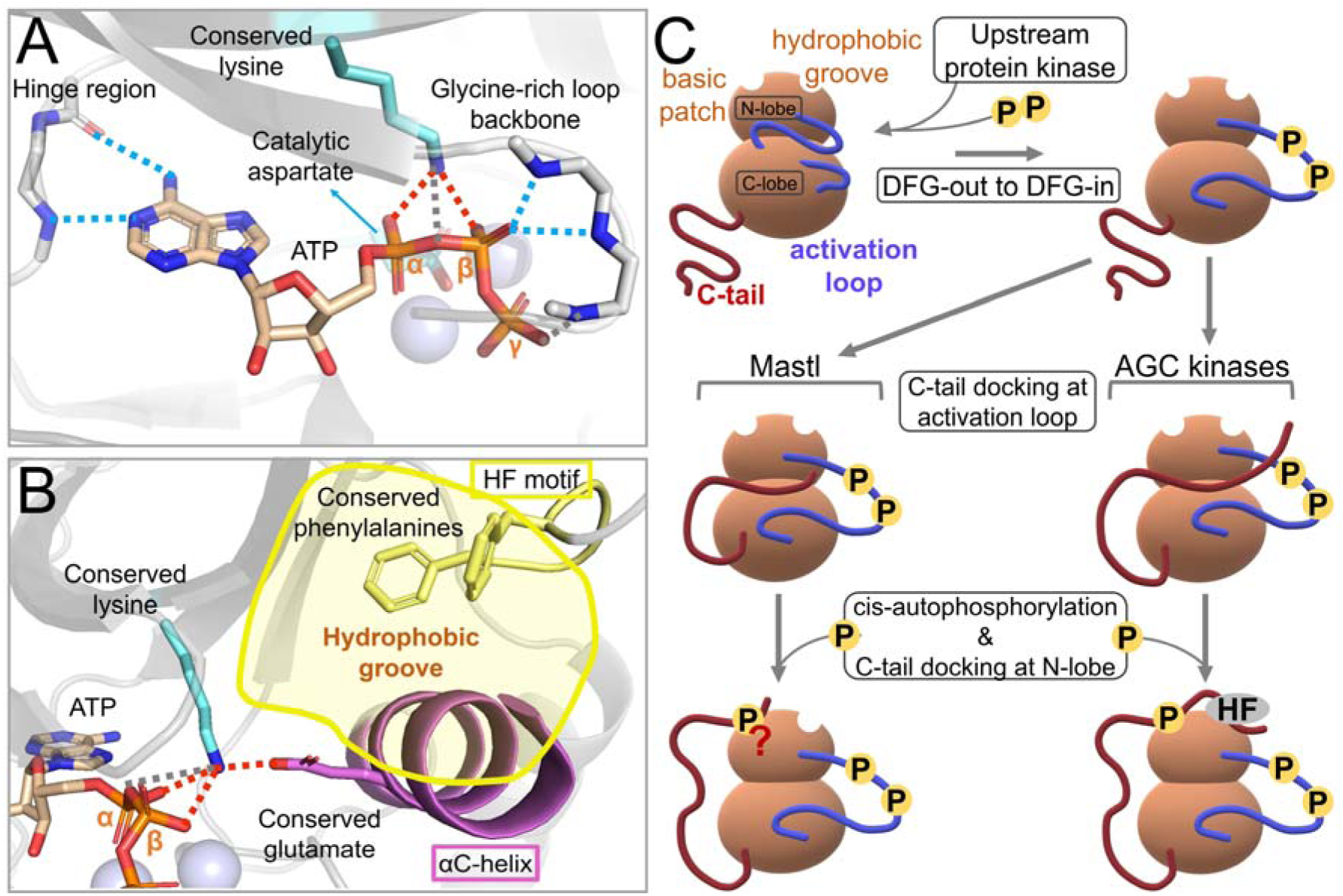
The structural features of AGC kinases. (A) The crystal structure of the catalytic subunit of ATP-bound c-AMP-dependent protein kinase (PDB entry 1ATP) (Zheng et al., 1993). The hinge region and glycine-rich loop backbones, catalytic aspartate, conserved lysine, and ATP are shown in stick forms. The manganese ions are depicted as transparent spheres. The observable ion-ion (salt bridge), ion-dipole, and dipole-dipole (hydrogen bonds only) interactions between the enzyme and the ATP are depicted as red, gray, and blue colored dashed lines, respectively. (B) The crystal structure of the catalytic subunit of ATP-bound c-AMP-dependent protein kinase (PDB entry 3X2W) (Das et al., 2015). The conserved lysine and glutamate, the conserved phenylalanine residues of the HF motif, and ATP are shown in stick forms. The αC-helix (violet), HF motif (yellow), and the proximal β trand (transparent gray) together form the hydrophobic groove. The hydrophobic groove is indicated by the transparent yellow area. The magnesium ions are depicted as transparent spheres. The observable ion-ion (salt bridge) and ion-dipole interactions between the enzyme and the ATP are depicted as red and gray colored dashed lines, respectively. The blue, orange, and red colored atoms are nitrogen, phosphorus, and oxygen, respectively. (C) Comparison of the general AGC kinase and Mastl kinase activation models. The bilobal protein kinase structure is depicted as light-brown bubbles. The white indents at the top-left and top-right positions of the N-lobe indicate the basic patch and the hydrophobic groove, respectively. The activation loop and the C-tail are depicted as blue and dark-red lines, respectively. In general, protein kinases are activated by phosphorylation of their activation loops. The cartoon illustrations show the difference between the activation of Mastl (left) and the general AGC kinase activation model (right). For simplicity, the extended NCMR is not represented in the illustrated Mastl kinase structures.

Activation loop starts with the DFG motif and typically encompasses the following 20-30 residues of the connecting loop (Modi & Dunbrack, 2019). This segment functions as a docking site for the substrate peptide. The sequence of the activation loop confers substrate specificity by determining the target consensus sequence through specific interactions. The conformation of the activation loop determines enzymatic activity by affecting three factors. These are accession of ATP to the ATP binding pocket, accession of the peptide substrate to the peptide binding groove, and orientation of the catalytic aspartate side chain. The active and inactive conformations of the activation loop are defined as DFG-in and DFG-out, respectively (Figure S1). The transition between DFG-in and DFG-out conformation is determined by the phosphorylation status of the activation loop. According to the general kinase activation model, DFG-in conformation is induced by activation loop phosphorylation. The phosphorylation event can be executed by upstream protein kinases or it can be autophosphorylation (Adams, 2003; Vijayan et al., 2015). In its inactive state, the activation loop simultaneously blocks its peptide substrate docking site and the ATP binding pocket. The catalytic aspartate side chain is tilted away from the ATP binding pocket. In its active state, activation loop simultaneously exposes its peptide substrate docking site and releases the ATP binding pocket. The catalytic aspartate side chain is tilted inwards the ATP binding pocket (Figure S1) (Huse & Kuriyan, 2002; Zhou et al., 2010).

In AGC family of protein kinases, phosphorylation of the activation loop can only partially activate the enzyme. A unique feature of the AGC kinases is that the core catalytic elements need to be stabilized through long-distance intramolecular interactions, in an allosteric manner. A series of conformational changes are induced by the C-tail/N-lobe interactions. The C-tail has two important regions that directly interact with the N-lobe. These are the turn motif and the HF motif (hydrophobic motif, FXXF). NLT (N-lobe tether) encompasses the stretch that proceeds the turn motif towards its C-terminal, including the HF motif. According to the currently accepted model, the conserved turn motif phosphosite aids docking of the C-tail to the basic patch at the N-lobe. Extending from this initial docking point via NLT, the HF motif reaches the hydrophobic groove of the N-lobe. The HF motif enhances the hydrophobic packing by inserting the benzene rings of its conserved phenylalanines into this hydrophobic groove. As a result, HF motif of the C-tail, αC-helix of the N-lobe, and the proximal β-strand of the N-lobe antiparallel β-sheet together construct a hydrophobic groove. This hydrophobic packing allows the C-tail to reposition the αC-helix. Ultimately, the conserved glutamate within the αC-helix and the conserved lysine located in the ATP binding pocket form a salt bridge. As explained previously, this conserved lysine functions in coordination of the ATP. Given the vital function of this lysine in the enzyme activity, it is clear that the C-tail indirectly aids the active site for coordination of the ATP. Briefly, through an intramolecular allosteric regulation, a conformational domino effect occurs. HF motif repositions the αC-helix, αC-helix coordinates the conserved lysine, and then this lysine ultimately contributes to coordination of the ATP (Figure 1B). Using the crystal structure of an atypical AGC kinase (Arencibia et al., 2017), human PKC iota (PDB entry 3A8W) (Takimura et al., 2010), the complete molecular machinery is visualized in Figure S2.

The C-tail can also exert its regulatory effects via other intramolecular interactions it establishes. It extends over the N-lobe to find the docking sites for the HF and the phosphorylated turn motifs. Doing so, the remaining N-terminal segment of the C-tail is vertically wrapped around the kinase. This wrapped middle segment engages in other intramolecular interactions (Kannan et al., 2007; Pearce et al., 2010; Taylor et al., 2012). Of note, PKA possesses a C-tail phosphate that does not contribute to intramolecular interactions (Figure S3), making the current model of AGC kinase activation questionable.

### 1.3. Activation of Mastl kinase

Mastl (Microtubule-associated serine/threonine kinase like) is a unique serine/threonine protein kinase among the AGC family kinases. It has an unusually long (∼500 residues) linker region between its N-terminal and C-terminal lobes. Studies on Xenopus egg extracts have revealed three phosphosites that are critical for the activation of Mastl (Blake-Hodek et al., 2012; Vigneron et al., 2011). Positions of these residues in mouse Mastl are T193, T206, and S861 (T194, T207, S875 in human; T193, T206, S883 in Xenopus). T193 and T206 are located in the activation loop within Cdk1 consensus motifs (proline-directed) and are phosphorylated by Cdk1 *in vitro*. These two phosphosites are necessary for nuclear export of Mastl before NEBD (nuclear envelope breakdown) (Alvarez-Fernandez et al., 2013). According to the currently accepted model, phosphorylation by Cdk1 partially activates Mastl. Activation loop phosphorylation corresponds to the initial step of general protein kinase activation. The activation loop shifts from DFG-out to DFG-in conformation. Upon this partial activation, Mastl undergoes cis-autophosphorylation by phosphorylating its turn motif (tail/linker) phosphosite S861 (Blake-Hodek et al., 2012). As explained in the previous section, in AGC kinases, this turn motif phosphoresidue is docked to a basic patch on the N-lobe, contributing to the tethering of the C-tail over N-lobe. In this regard, the turn motif phosphorylation serves as a means to guide the HF motif towards the hydrophobic groove. However, in the case of Mastl, the C-tail of the enzyme is shorter and it does not possess a known HF motif. Therefore, it cannot reach the hydrophobic groove (Figure 1B).

Results of Vigneron et al. (2011) demonstrated that Mastl has to be partially active in order for its C-tail phosphosite to undergo autophosphorylation. Later, Blake-Hodek et al. (2012) showed that the C-tail autophosphorylation is an intramolecular reaction, i.e., cis-autophosphorylation. After C-tail phosphorylation, Mastl undergoes C-tail/N-lobe docking, suggested by the alanine scanning mutagenesis study of Vigneron et al. (2011). In their experiment, they targeted the C-tail phosphosite and its proximal lysine residues of N-lobe. They have also observed that activity of constitutively active Mastl increases up to 156% in the presence of a synthetic peptide (PIFtide) that mimics the hydrophobic motif. Their finding suggests that Mastl utilizes the HF motif of other AGC kinases in a heteromeric complex to complete its activation. However, a decade later, it is still not clear what purpose would C-tail/N-lobe intramolecular docking would serve, when the C-tail is devoid of HF motif. This raises the question whether the C-tail phosphorylation is not essential for Mastl activation.

The phosphorylation-dependent activation mechanism of Mastl and the biological significance of its NCMR have been extensively studied *in vitro* and using Xenopus oocyte extracts (Blake-Hodek et al., 2012; Vigneron et al., 2011). According to the current model, phosphorylation of the two activation loop phosphosites (T193 and T206 in mouse) and the C-tail turn motif phosphosite (S861 in mouse) are crucial for the activation of Mastl. In the present study, we have reevaluated the biological significance of these three phosphosites and two additional phosphosites (S212 and T727) that were proposed to be potential contributors to Mastl activation (Blake-Hodek et al., 2012). Other than these, we tested the hyperactive Drosophila *Scant* mutation K71M (Archambault et al., 2007), thrombocytopenia-associated mutation E166D (Hurtado et al., 2018), and two different NCMR deletions (Δ194-725 and Δ305-620). We obtained viable Mastl knockout MEF clones that express the ectopic E166D, K71M, S861A, or S861D mutants. Given the vital role of the turn motif phosphosite S861 in AGC kinase activation, we focused our study on the functional characterization of this phosphosite.

## 2. METHODS

### 2.1. Software

CLC Main Workbench 7.9.1 software was used for sequence visualization and analysis (https://digitalinsights.qiagen.com). The gel images, immunoblot images, and micrographs were processed using Adobe Photoshop and GIMP (version 2.10.12) (The GIMP Development Team, 2019). The boxplots were drawn in R (v3.6.1) (R Core Team, 2019), invoked by RStudio (v1.2.5001) (RStudio Team, 2015), using the ggplot2 (Kassambara, 2019) R package.

### 2.2. Structural modeling of Mastl

The crystal structure coordinate files were obtained from the Protein Data Bank (PDB) (https://www.rcsb.org/) (Berman et al., 2000). The coordinate files were visualized by using PyMOL (version 2.4.0a0) (Schrödinger, LLC, 2015). By the help of the sequence alignments given in one of the reference studies (Blake-Hodek et al., 2012), the domain and motif boundaries of mouse Mastl were determined by using the UniProt database (https://www.uniprot.org/) (The UniProt Consortium, 2019) and EMBOSS Needle webserver (https://www.ebi.ac.uk/Tools/psa/emboss_needle/) (Rice et al., 2000). Briefly, the FASTA format sequences were obtained from UniProt and the pairwise sequence alignments were performed using EMBOSS Needle. Based on the determined motif boundaries, the motifs were mapped on the 3D structure visuals for figures. Mouse Mastl was modeled by MODELLER 9.25 (https://salilab.org/modeller/) (Fiser et al., 2000; Martí-Renom et al., 2000; Šali & Blundell, 1993; Webb & Sali, 2016), invoked by MODOMAX (https://github.com/MehmetErguven/MODOMAX) (Erguven & Karaca, 2021).

Structural refinements, binding energy, and buried surface area calculations were performed using the Guru interface of HADDOCK2.2 web server (https://milou.science.uu.nl/services/HADDOCK2.2/haddockserver-guru.html) (van Zundert et al., 2016). When the docking stages are skipped, HADDOCK2.2 is capable of refining the interaction of separate molecules. To refine the C-tail and N-lobe interactions, the protein kinase chain was separated by the N-terminal starting position of C-tail. The C-tail and the rest of the protein were submitted as if they were separate chains.

The following changes were applied to the default docking protocol;

- “Remove non-polar hydrogens?” checkbox was unchecked under the “Distance restraints” tab.
- “Define center of mass restraints to enforce contact between the molecules” checkbox was checked under the “Distance restraints” tab.
- “Randomize starting orientations”, “Perform initial rigid body minimization”, and “Allow translation in rigid body minimization” checkboxes were unchecked under the “Advanced sampling parameters” tab.
- “number of MD steps for rigid body high temperature TAD”, “number of MD steps during first rigid body cooling stage”, “number of MD steps during second cooling stage with flexible side-chains at interface”, and “number of MD steps during third cooling stage with fully flexible interface” parameters were set to 0 under the “Advanced sampling parameters” tab.

HADDOCK outputs residue-based energy contributions of each interfacial amino acid (expressed in electrostatics, vdW and electrostatics+vdW energy terms), deposited in *ene-residue.disp* file (located under HADDOCK output folder: structures/it1/water/analysis). For each interface residue, various types of scores were extracted from the HADDOCK runs for its 200 different states. These scores were presented in boxplot format in which the energy distribution of the residues can be compared to one another (Figure 6, S7).

Protein surface electrostatics potentials were calculated using the APBS- PDB2PQR software suite web server (https://server.poissonboltzmann.org/) (Baker et al., 2001; Dolinsky et al., 2004).

### 2.3. Molecular cloning

N-terminally HA-tagged, retroviral expression constructs (pBABE-Puro) for mouse Mastl WT, D155N (kinase-dead, KD), E166D (thrombocytopenia-associated mutant), and K71M (Drosophila *Scant* mutation) were generated in Kaldis Laboratory (Institute of Molecular and Cell Biology, Singapore). The other mutations were created by site-directed mutagenesis. The primers used for site-directed mutagenesis are given in Table S2.

### 2.4. PCR genotyping

A three-primer strategy was utilized for genotyping of Mastl locus, to distinguish FLOX and KO alleles (Figure 2A). The primers used for genotyping; are given in Table S2. For the PCR genotyping, the samples were prepared according to the adapted HotSHOT protocol (Truett et al., 2000). For the genotyping PCR, 0.5 µL of DNA sample and 0.25 units of Taq DNA polymerase (A111103; Ampliqon) was used for each 10 µL of reaction volume. Final primer concentrations were 1 µM for FOR and REV1, and 0.15 µM for REV2 primers. 35 PCR cycles were performed using 67°C annealing temperature. Under these optimized conditions, the PCR reaction consistently yields roughly equal intensity bands for Mastl^FLOX^ (300 bp) and Mastl^KO^ (500 bp) alleles from a heterozygous Mastl^FLOX/KO^ cell clone (Figure 2).

**Figure 2.**
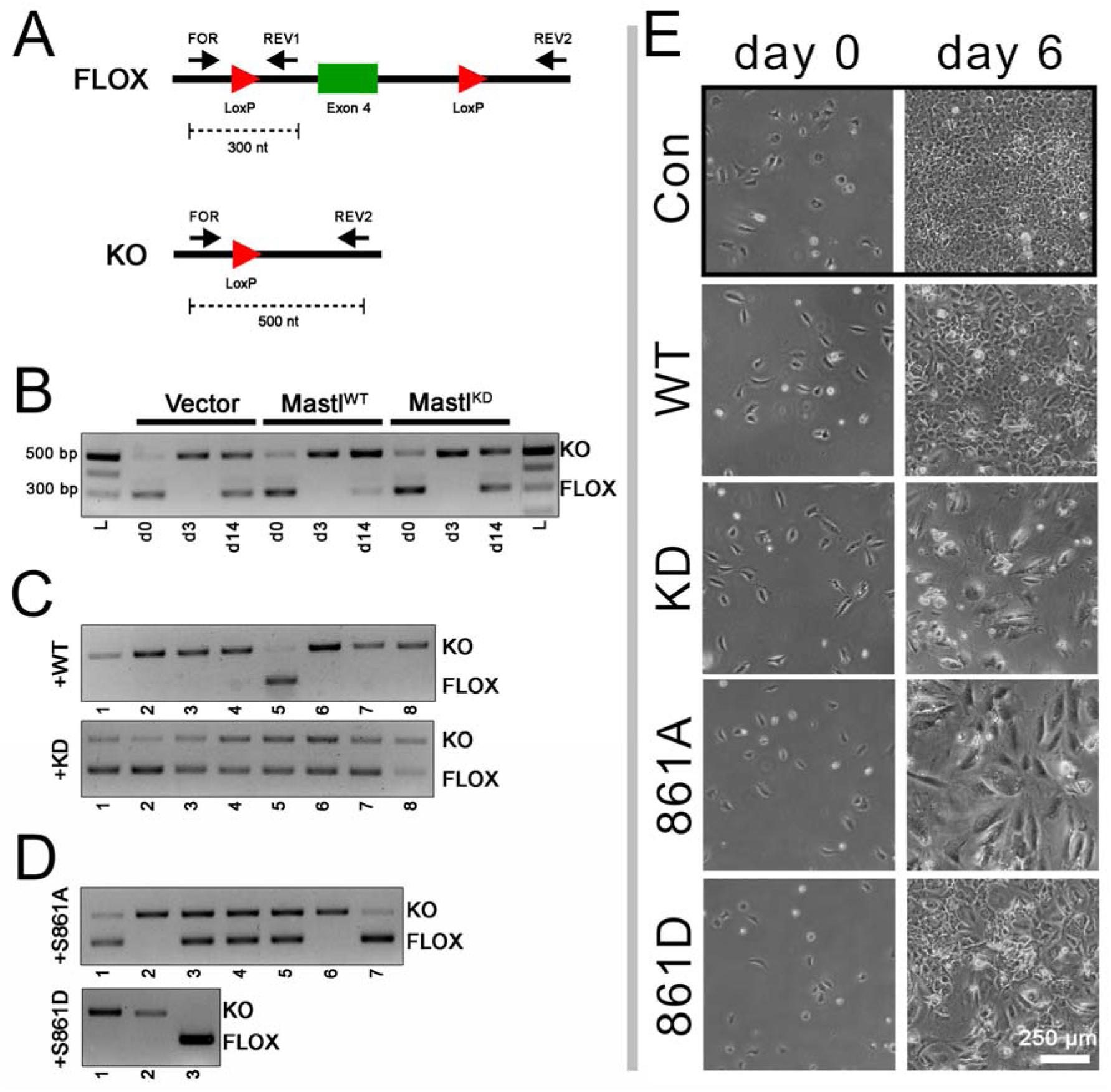
Genotype and phenotype analysis of the ectopic Mastl-expressing Mastl conditional knockout (cKO) MEFs. (A) A three-primer strategy was employed for PCR genotyping of the Mastl locus. PCR amplification yields 300 and 500 bp fragments for Flox and KO alleles respectively, at roughly equal intensities from a heterozygous Mastl^Flox/KO^ clone’s DNA. (B) Stable cell pools were treated with 4-OHT and cells were genotyped at the indicated days. Three days after 4-OHT addition, FLOX allele is undetectable. However, after 14 days, it becomes visible again, and especially prominent in stable cell pools infected with empty vector and Mastl^KD^ construct. (C) After limited dilution, clonal cell lines were isolated and genotyped. Mastl^WT^ construct can complement endogenous Mastl loss. Hence, in majority of the clones, endogenous Mastl is deleted. None of the Mastl^KD^ infected clones have lost both endogenous Mastl loci. (D) From Mastl^861A^ and Mastl^861D^ infected pools, two clones could be isolated for both. (E) Proliferation of stable cell pools after deletion of endogenous Mastl. The micrographs were acquired immediately before (day 0) and six days (day 6) after 4-OHT treatment. Con indicates untreated cells. Scale bar is 250 μm.

### 2.5. Cell culture, retroviral transduction, and stable cell line engineering

The PLAT-E cells (Cell Biolabs) and Mastl conditional knockout immortalized MEFs (Diril et al., 2016) were grown in DMEM supplemented with 10% FBS, and antibiotics. Cells were cultured in a 95% humidified incubator with 5% CO_2_ at 37°C. Lipofectamine 3000 transfection reagent (L3000015; Thermo Scientific) was used for transfection of PLAT-E cells. PLAT-E culture and virus production were carried out according to the manufacturer’s instructions. The viral supernatant was harvested 2 days and 3 days post-transfection, and stored at 4°C up to a week until use.

Polybrene was mixed with retroviral medium to 8 µg/mL final concentration, immediately before infection of MEFs. MEFs were transduced at 40% confluence. Viral media collected at the second day of transfection was used in the first round of transduction. One day post-transduction, the used viral media was removed and a second round of transduction was performed by using the viral media that was harvested at the third day of transfection. 8 hours after the second round of transduction, the viral media was removed. Cells were cultured further in normal growth medium overnight for recovery, prior to antibiotic selection. The stable cell lines were generated under 2-4 µg/mL puromycin selection. Cells were selected for a week and negative control cells died within the first two days of selection. Mastl knockout was induced by adding 4-OHT to the culture medium at 20 ng/mL final concentration. The control cells were treated with an equal volume of DMSO. The knockout induced stable cell lines were subjected to limited dilution approximately 6 days post-induction.

### 2.6. Western blot and immunocytochemistry

Commercially available primary antibodies used for western blot are rat anti-HA tag (clone 3F10; Roche), rabbit anti-Cyclin B1 (4138S; Cell Signaling Technology), mouse anti-Cdk1 (sc-54; Santa Cruz Biotechnology), and mouse anti-HSP90 (610419; BD Biosciences). The rabbit anti-Mastl polyclonal antibody was self made in Kaldis laboratory (Diril et al., 2016).

### 2.7. Cell viability and proliferation assays

The alamarBlue proliferation assay was carried out in 96-well plate format in three replicates. The daily measurements were initiated one day after seeding the cells. Cells were incubated in 150 µL of assay medium for 4 hours. The assay medium was prepared by diluting 1 volume of alamarBlue dye reagent (BUF012A; Bio-Rad) in 9 volumes of growth medium. The metabolic activity was quantified fluorometrically by using 560 nm excitation wavelength and recording the emission at 590 nm. The assay was performed for 6 successive days.

The proliferation rate was also measured by a modified 3T3 assay (Diril et al., 2012) in 6-well plates. 20000 cells were seeded per well and cells were counted after 3 days and 20000 cells were plated again. Counts were repeated for four successive passages.

## 3. RESULTS AND DISCUSSION

### 3.1. A robust rescue-assay for evaluation of the Mastl mutants

We used the following workflow in order to assess the biological significance of the previously described mutations and NCMR deletions (Blake-Hodek et al., 2012; Vigneron et al., 2011). Briefly, (i) conditional knockout (cKO) Mastl^FLOX/FLOX^ cell pools stably expressing Mastl mutants were generated, (ii) endogenous Mastl gene was deleted, (iii) clonal cell lines were isolated and genotyped, and (iv) the clones whose endogenous Mastl genes were knocked out were further characterized.

The phosphosites of interest that we screened are T193, T206, S212, T727, and S861. For the phosphosite studies, we have used the T193V, T206V, S212A, T727V, S861A non-phosphorylatable mutants and the T193E, T206E, S861D phosphomimetic mutants. Other Mastl variants in the present study are E166D (thrombocytopenia associated variant) (Hurtado et al., 2018) and K71M (*Scant*) (Archambault et al., 2007) point mutations, and partial (Δ305-620) and full (Δ194-725) NCMR deletions (Blake-Hodek et al., 2012). KD (kinase-dead) variant and WT (wild-type) Mastl were used as negative and positive controls, respectively.

MEFs were infected with retroviral expression constructs (pBABE-Puro) and selected with puromycin, in order to establish stable cell pools expressing ectopic Mastl mutants. To quickly assess the complementation capacity of the ectopic constructs, the stable pools were treated with 4-OHT, and the cells collected before and after KO induction (3 and 14 days) were genotyped (Figure 2A and 2B). Due to the nature of the inducible Cre systems, cKO MEFs include a basal level of knockout cells/loci as a result of spontaneous Cre leakage into the nucleus, even without 4-OHT induction (Zhong et al.). Three days post-induction, a complete knockout of the Mastl gene was observed. However, within fourteen days, initially undetectable levels of Mastl^FLOX/FLOX^ or Mastl^FLOX/KO^ cells, expressing endogenous Mastl, proliferate. Cells escaping knockout is a known disadvantage of the inducible Cre systems (Diril et al. 2012). Cells expressing endogenous Mastl account for the majority of the cell population when the ectopic constructs cannot complement endogenous Mastl loss (see day 14 in Figure 2B).

Although, nonfunctional Mastl (i.e., Mastl^KD^) expressing cell pools eventually proliferate after 4-OHT treatment, there is a lag compared to the functional Mastl (i.e., Mastl^WT^) expressing cells. Analysis of different Mastl mutant cell pools six days after 4-OHT treatment suggests that S861A, S861D, K71M, and E166D mutations retain partial or full kinase activity compared to WT (Figure 2E and S5). These results are consistent with the genotyping analysis (Figure 2D and S4A) which confirms that cells expressing these mutants are viable when their endogenous Mastl is knocked out.

Clonal cell lines were generated after 4-OHT treatment of cell pools expressing various Mastl mutants. PCR genotyping of WT and KD expressing pools confirmed that our experimental system works. As expected, multiple endogenous Mastl KO clones could be isolated from the pool expressing ectopic WT Mastl, whereas none from the KD Mastl-expressing pool (Figure 2C). Analysis of K71M (Drosophila *scant*) and E166D (thrombocytopenia associated mutation) clones showed that these Mastl mutants can also complement endogenous Mastl loss (Figure S4 and S5). This is an expected result as these mutations do not have a significant effect on cellular proliferation (Archambault et al., 2007; Hurtado et al., 2018). None of the NCMR deletion clones had homozygous KO genotype, suggesting that the deletions cover catalytic or structural elements that are essential for Mastl function *in vivo*. This is in line with previous observations for NCMR deletion mutants (Vigneron et al., 2011). On the other hand, the number of clones we genotyped for our deletion mutants were limited. Therefore, we cannot rule out the fact that with an increased sample size, a homozygous KO clone may be obtained.

After establishing the complementation screening strategy and successfully testing it on known viable mutations (K71M and E166D), we next focused on Mastl phosphorylation sites with putative roles on kinase activation. We created constructs with point mutations that have non-phosphorylatable (T193V, T206V, S212A, T727V, and S861A) or phosphomimetic residues (T193E, T206E, and S861D). We were able to isolate clonal cell lines for S861A and S861D, in which, the endogenous Mastl gene was knocked out (Figure 2D). Other mutations did not yield viable cell lines in our screen, as all the clones expressed one or both copies of the endogenous Mastl (Figure S4).

### 3.2. The turn motif phosphosite mutant Mastl is viable

Our complementation screening showed that the turn motif phosphosite mutant expressing clones (S861A and S861D) were viable even if the endogenous Mastl gene was deleted. (Figure 2A). As previously explained, the turn motif phosphorylation is known to be crucial for AGC kinase activation. Therefore, we focused our study on these knockout clones for functional characterization of this particular phosphosite. We selected one clone from each of the WT, S861A, and S861D clonal cell lines for further experiments.

Western blot analysis of the selected clones confirmed the expression of the ectopic constructs (Figure 3A). Although the expression levels of the ectopic proteins were lower than the endogenous Mastl levels, this was sufficient for cellular proliferation. Especially S861D clone, whose Mastl protein level was significantly low, could still proliferate albeit at a slightly reduced rate (Figure 4).

**Figure 3.**
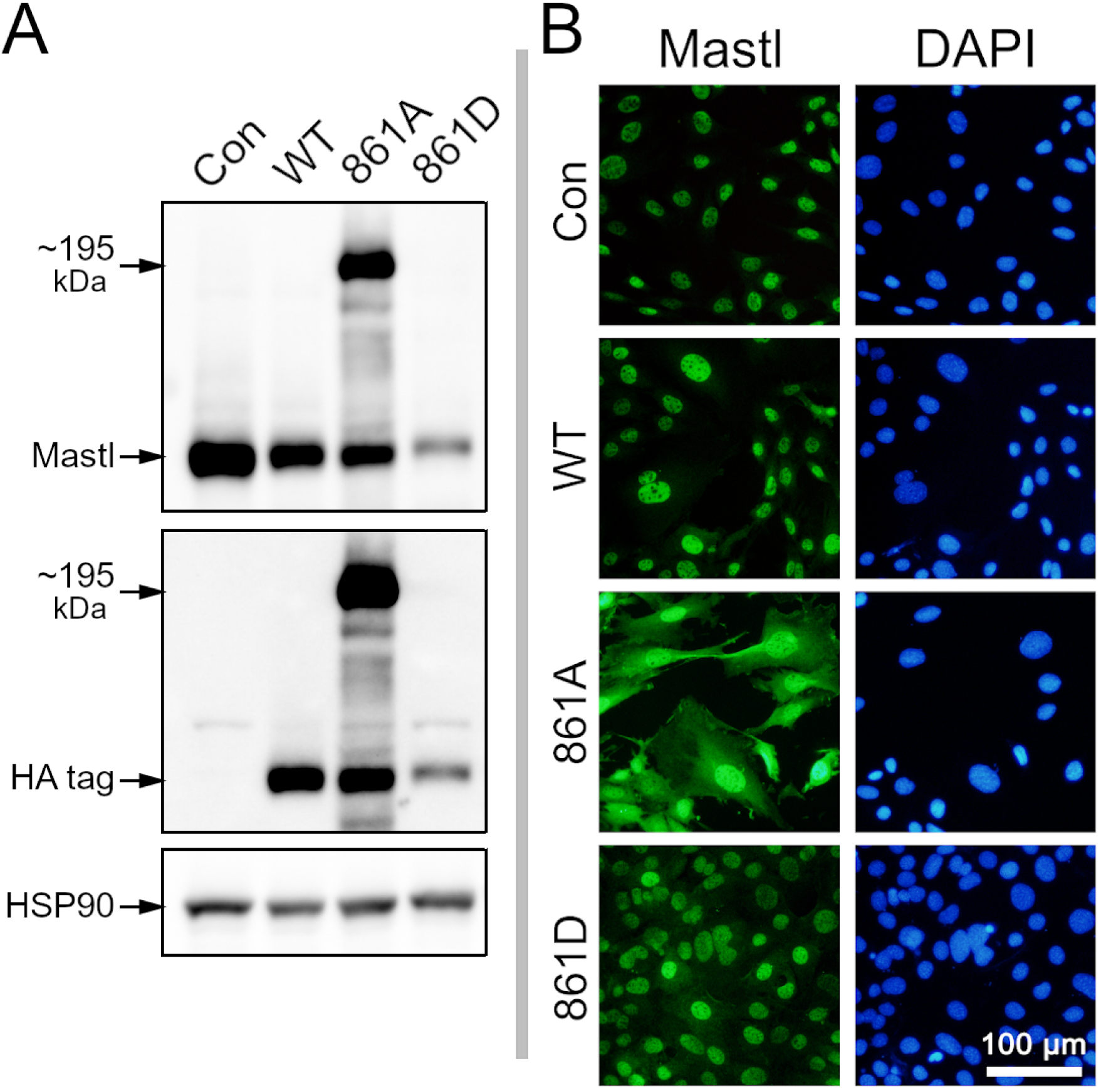
Protein analysis of the ectopic Mastl-expressing homozygous knockout clonal cell lines. (A) Western blot analysis of ectopic Mastl expression. Anti-Mastl and anti-HA antibodies were used to detect and compare Mastl levels in control (Mastl locus intact) and ectopic Mastl expressing clones. HSP90 was used as loading control. The selected S861A clone had an additional higher molecular weight Mastl product that runs around 195 kDa. This was not due to the plasmid vector we created but possibly results from in-frame integration of the ectopic Mastl coding sequence into another gene. Additional clone screening identified S861A clones with expected Mastl MW (data not shown). (B) Analysis of the ectopic Mastl localization. Cells were fixed and stained with anti-Mastl antibodies. Ectopic Mastl protein is (green) localized to the nucleus (blue, DAPI). S861A clone that expressed a high molecular weight fusion product displays additional cytosolic distribution. Scale bar is 100 μm.

**Figure 4.**
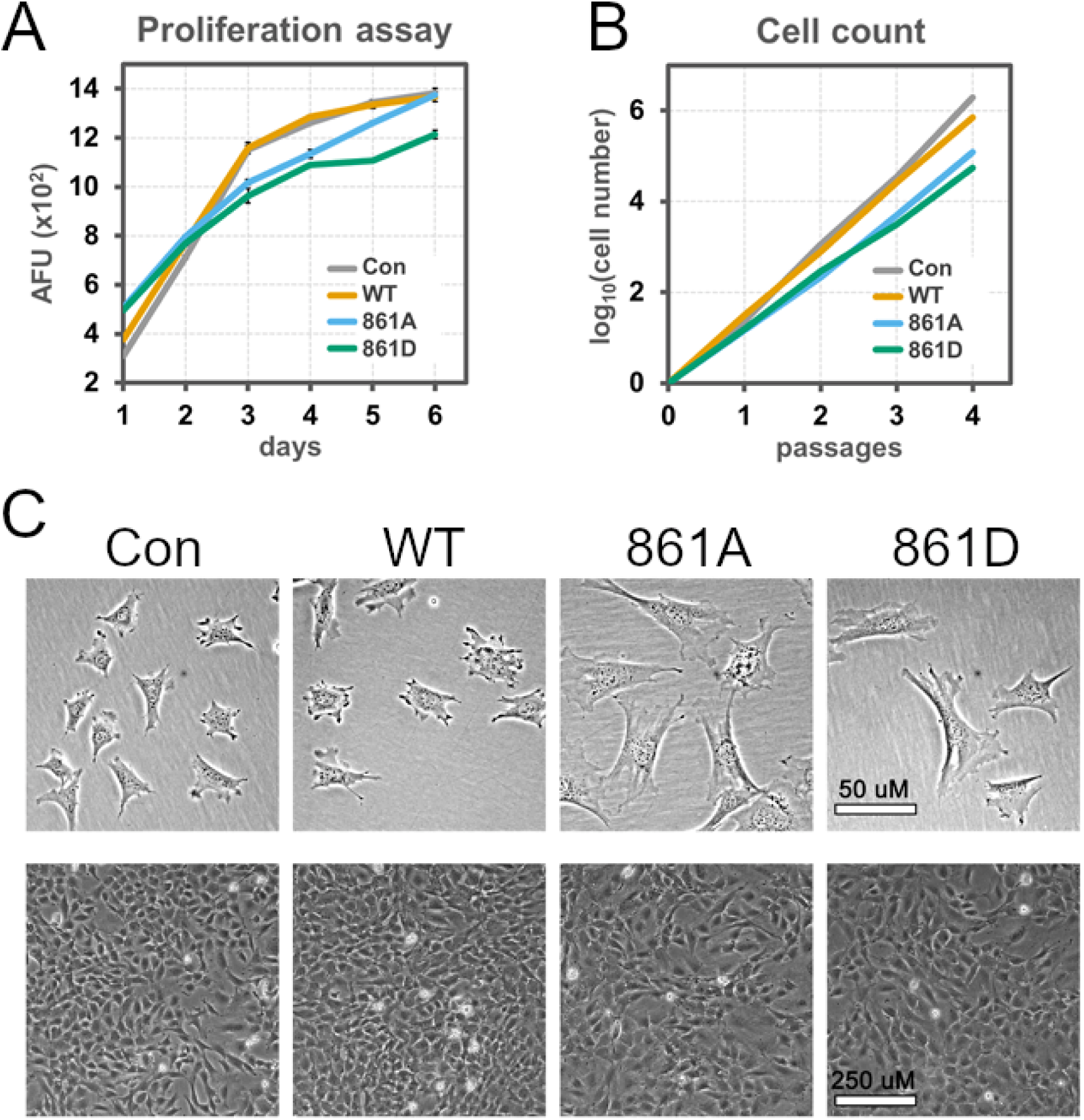
Proliferation rates of control, WT, 861A and 861D clones (A) alamarBlue proliferation assay of the control MEFs and clonal cell lines. The metabolic was fluorometrically measured (AFU: arbitrary fluorescence units) in terms of the resazurin reduction activity. The cells were monitored for six days. (B) A modified 3T3 assay was performed to directly measure the increase in cell number over time. Number of cells at the beginning of the assay were normalized to one. (C) For 3T3 assay, the cells were seeded at low density (20000 cells/well in 6-well plates) in order to show their unique morphologies in the absence of cell-cell contact restrictions (upper panel). Three days after the seeding, all clones become confluent (lower panel). Scale bar is 50 µm in upper panel and 250 µm in lower.

Subcellular localization of the ectopic Mastl proteins were analyzed by immunocytochemistry staining. As expected, endogenous and ectopic Mastl WT proteins localized to the nucleus. Mastl S861A and S861D mutants also localized to the nucleus (Figure 3B). Therefore, S861 phosphorylation has no effect on Mastl localization.

### 3.3. The Mastl S861 phosphosite mutation does not impair cell proliferation

We have demonstrated that deletion of endogenous Mastl can be rescued when Mastl S861A or Mastl S861D mutants are expressed ectopically. To compare the proliferation rates of these clones to that of control MEFs or Mastl WT-expressing clones, we decided to undertake quantifiable assays.

First, the proliferation rates of these clones were measured indirectly by alamarBlue proliferation assay which measures the overall metabolic activity. The results suggest that the mutant Mastl expressing clonal cell lines have proliferation rates comparable to control cells or Mastl WT expressing clonal cell line (Figure 4A). Next, we measured the proliferation rates directly by a modified 3T3 assay (see Methods). The mutant clones had slightly reduced proliferation rates compared to control cells or the WT clone (Figure 4B, C).

These results suggest that, S861 phosphorylation site mutations do not have a remarkable negative effect on cell proliferation rate. In fact, the slight reduction in proliferation rates of the mutant clones is possibly due to the lower expression level of ectopic Mastl proteins (See Figure 3A).

### 3.4. Mammalian Mastl kinase is regulated by cis-autophosphorylation

Cis-autophosphorylation of Mastl kinase has been previously shown in Xenopus egg extracts. WT Mastl kinase purified from okadaic acid treated Sf9 cells was capable of phosphorylating itself, whereas its kinase-dead mutant was not. When WT active Mastl was added to an excess of inactive Mastl, trace amounts of trans-autophosphorylation of the inactive Mastl was observable. Cis-autophosphorylation of the active WT Mastl was by far the most abundant event (Blake-Hodek et al., 2012). This indicates that the autophosphorylation event is dominated by cis-autophosphorylation.

In the present study, we have validated the cis-autophosphorylation model using intact MEFs. In mitotically arrested primary MEFs, a slower running Mastl isoform appears in immunoblots. The high molecular weight band is due to the decreased electrophoretic mobility of the hyperphosphorylated Mastl protein, and it can be reversed by phosphatase treatment of the protein extract (Figure 5A, B). In fact, mitotic cells only have the hyperphosphorylated Mastl isoform, whereas it is undetectable in interphasic cells (Figure 5C).

**Figure 5.**
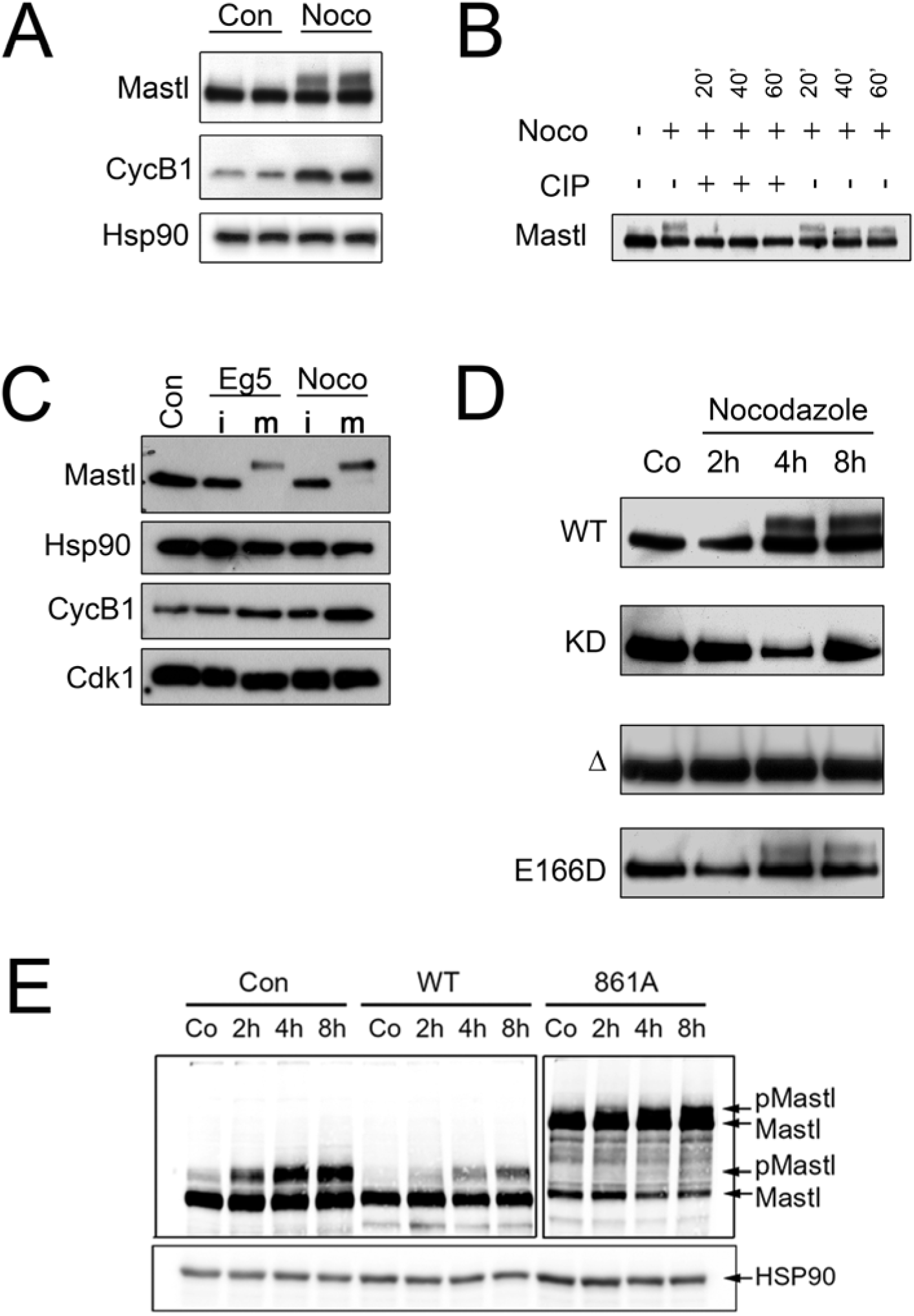
Mastl undergoes cis-autophosphorylation. (A) Mitotic arrest of asynchronously proliferating primary MEFs results in phosphorylation of Mastl which can be detected by a higher MW band in western blots. Cyclin B1 was used an indicator of mitotic arrest and two biological replicates were analyzed. (B) CIP (calf intestinal phosphatase) treatment of the protein extracts from nocodazole-arrested cells results in disappearance of the putative phosphorylated Mastl isoform. (C) Primary MEFs were treated with nocodazole or Eg5 to enrich mitotic arrested cells. Mitotic cells (m) were separated from interphase cells (i) by mitotic shake off and protein extracts were analyzed by WB. Phosphorylated and non-phosphorylated Mastl are exclusively present in mitotic and interphase cells respectively. (D) Wildtype MEFs expressing ectopic WT, KD, NCMR deletion, and E166D Mastl variants (HA-tagged) were treated with nocodazole to enrich mitotic cells. WT and E166D variants accumulated a high MW phosphorylated Mastl band. However, KD or NCMR deletion variants do not undergo a size shift indicating that they do not acquire phosphorylations that can decrease their electrophoretic mobility. Therefore, phosphorylation-dependent size shift is due to cis-autophosphorylation. Anti-HA antibodies were used to detect the exogenous Mastl variants. (E) Clonal cells lines expressing ectopic WT or S861A Mastl variant were arrested in mitosis and analyzed by WB. S861A mutation does not hinder cis-autophosphorylation. The blots were immunostained with anti-Mastl antibody.

To investigate how the hyperphosphorylation takes place, we ectopically expressed the WT or kinase-dead HA-tagged mouse Mastl in WT MEFs. The cells were treated with nocodazole for different time intervals to enrich the mitotic cells. Nocodazole treatment results in accumulation of the phosphorylated mitotic proteins due to metaphase arrest of the cells. Consequently, the proportion of phosphorylated Mastl increases (Diril et al., 2016). The hyperphosphorylated Mastl has a decreased electrophoretic mobility, which is observed as a band upshift in the immunoblots. The Western blots against the HA-tagged ectopic proteins show that the MEFs expressing WT HA-Mastl display the upshift, whereas MEFs expressing the kinase-dead mutant do not (Figure 5D). Since these cells do not rely on the ectopic kinase for mitotic entry as they express endogenous WT Mastl, the presence of the upshift only in the WT or viable mutants of Mastl (E166D) can best be explained by cis-autophosphorylation. Trans-autophosphorylation or phosphorylation by other kinases does not result in hyperphosphorylation. In contrast to earlier published reports (Blake-Hodek et al., 2012), we could not observe traces of trans-autophosphorylation. A possible explanation for this can be that the different protein detection techniques may have different limits of detection.

Expanding on this, we investigated whether Mastl S861A can become hyperphosphorylated. To that end, we repeated the mitotic arrest and autophosphorylation analysis for this clone.

It is possible that the ∼195 kDa fusion Mastl product (that is probably a result of an in-frame genomic integration) in S861A clone is not stable. This could explain its degradation products appearing as a smear in the blot. For the S861A clone, we observed trace amounts of upshift of the normal molecular weight Mastl. However, this upshift is masked by the said degradation products of the larger (∼195 kDa) Mastl. On the other hand, a shorter-distance, but denser upshift is observed for the ∼195 kDa Mastl S861A product (Figure 5E), which suggests it is also catalytically active. Based on this, it can be hypothesized that Mastl S861A is capable of cis-autophosphorylation.

### 3.5. Positioning of the phosphosite defines the C-tail conformation

Our experimental results suggest that pS861 phosphoresidue is auxiliary for Mastl kinase activation. To mechanistically explain this result, we decided to analyze the C-tail intramolecular interactions of Mastl and other AGC kinases. Currently, there is only one crystal structure of Mastl (*Homo sapiens*, 5LOH) available in the PDB. However, the structure is a truncated form of the enzyme, lacking the C-tail. Because of this, we decided to find suitable templates for homology modeling of Mastl.

For this, we have collected all available AGC kinase crystal structures that possess a phosphorylated C-tail. According to this criterion, 17 structures of two different AGC kinases are available in PDB, which are PKC iota and PKA (Table S1). The structures fall into two distinct groups based on the conformation of their phosphorylated C-tail residue. The phosphoresidue is either buried in the N-lobe, or it is solvent-exposed (Figure S3). Upon observing that there are two distinct C-tail phosphoresidue conformations, we first decided to elucidate which conformation would Mastl pS861 possess (buried or solvent-exposed). To this end, we performed pairwise sequence alignments between the kinase domains of mouse Mastl and human PKC iota or mouse PKA (Figure S6).

The sequence alignments reveal that mouse Mastl C-tail phosphosite does not correspond to a phosphoresidue in PKA and PKC iota, but it corresponds to an aspartate in PKC iota. To get a detailed insight, we obtained two separate homology models for Mastl, by using PKC iota (3A8W) or PKA (3FJQ) as the templates. Expectedly, the C-tail phosphosite of the model structures possesses the distinct conformation (i.e., solvent-exposed or buried) of the corresponding residue in their respective templates (Figure 6). Using HADDOCK2.2 web server, the binding score of the C-tail to N-lobe was predicted by using van der Waals score, electrostatics score, desolvation, and the combined HADDOCK scoring function (Figure S7). The binding score was calculated on the nonphosphorylatable, wild type, phosphomimetic, and phosphorylated forms of PDB 3A8W (PKC iota), PBD 3FJQ (PKA), 3A8W-based Mastl model, and 3FJQ-based Mastl model. Due to the charged interactions between the phosphoresidue and basic residues, we have turned our attention to the electrostatics scores of the interfacial residues. As expectedly, when the C-tail phosphoresidue is buried inwards the N-lobe, its contribution to the binding energies dramatically increases (Figure 6).

**Figure 6.**
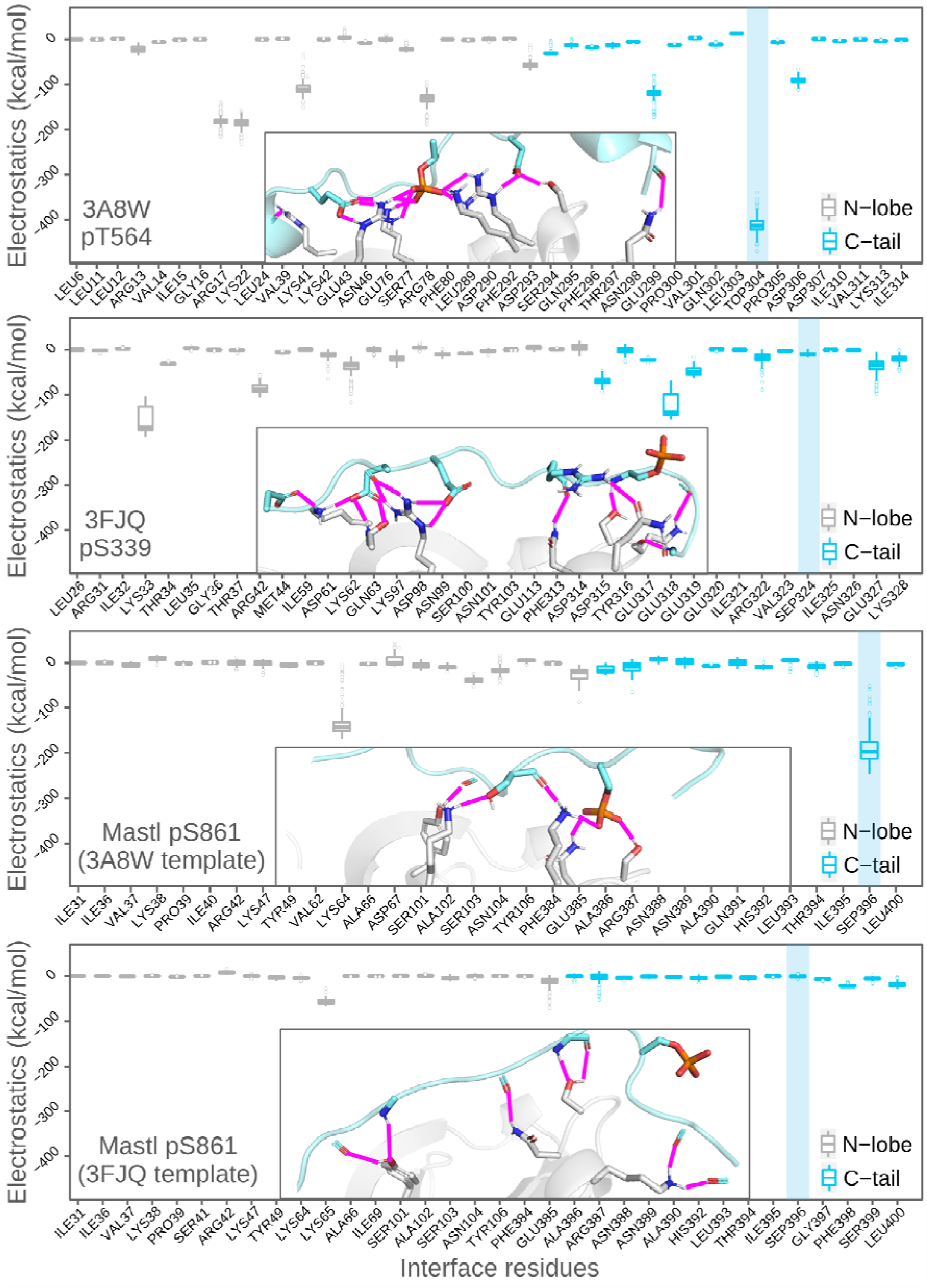
The PKC iota crystal structure (3A8W), PKA crystal structure (3FJQ), mouse Mastl 3A8W-based model structure, and mouse Mastl 3FJQ-based model structure were subjected to structural refinement using HADDOCK2.2 web server. For each structure, simulation results of their phosphorylated forms are given. Throughout the energy minimization, HADDOCK generated 200 states for each starting structure. For each interface residue, electrostatics scores of these 200 model structures are given as boxplots. Each box shows the energy distribution of individual residues throughout the simulation. For each run, among the 200 generated model structures, the one that has the lowest (best) HADDOCK score was selected for three-dimensional depiction of the interfaces, given in the respective plot areas. Both in plot and in the structure view, the C-tail turn motif and the N-lobe are cyan and gray colored, respectively. The transparent blue rectangle on the right side of each plot indicates the position of the phosphosite. The interfacial polar contacts were detected by PyMOL. The residues that form polar contacts are depicted as sticks. The polar contacts are depicted as magenta lines. The blue, orange, and red colored atoms are nitrogen, phosphorus, and oxygen, respectively. The residues are numbered according to their positions within the truncated structures. The positions of the phosphosites within the full-length sequences are written in the bottom-left corner of each plot.

Our computational results demonstrate that when the phosphoresidue is buried, it greatly contributes to the intramolecular interactions and when it is solvent-exposed, the phosphoresidue does not contribute to the C-tail docking. The pattern of polar contact distribution shows that in PKC iota and PKC-iota-based Mastl model, the contacts are clustered at the center, whereas in case of PKA and PKA-based Mastl model, contacts are clustered more at the edges of the interface (Figure 6). This two-sided stitching of the PKA C-tail might be a structural feature that has evolved as a means to promote phosphorylation-independent docking. We then evaluated the structural impacts of the nonphosphorylatable or phosphomimetic mutations of the C-tail phosphosite. To this end, we selected the HADDOCK-refined structures with the lowest HADDOCK score for the said mutants of PKC iota (based on PDB entry 3A8W) and PKA (based on PDB entry 3FJQ) in order to use for APBS (Adaptive Poisson-Boltzmann Solver) calculations. The surface electrostatics potentials of the proteins were calculated and mapped onto the three-dimensional structures (Figure 7). The electrostatics potential maps show that phosphomimetic glutamate mutation in PKC iota is interacting with the basic patch (the blue cavity). When the C-tail phosphosite is mutated to the nonphosphorylatable valine, the basic patch is expanded. In other words, the distance between the edges of the basic patch increases. This is because of the fact that, in the absence of a negative charge where there should be the phosphoresidue, the basic patch residues cannot engage in a polar contact network. On the other hand, in case of PKA, the phosphomimetic or nonphosphorylatable mutations of the C-tail phosphosite do not cause a remarkable structural difference. This is because the C-tail phosphosite is already tilted away from the N-lobe (Figure 7). These results together reinforce our hypothesis that Mastl pS861 phosphoresidue does not contribute to intramolecular C-tail docking.

**Figure 7.**
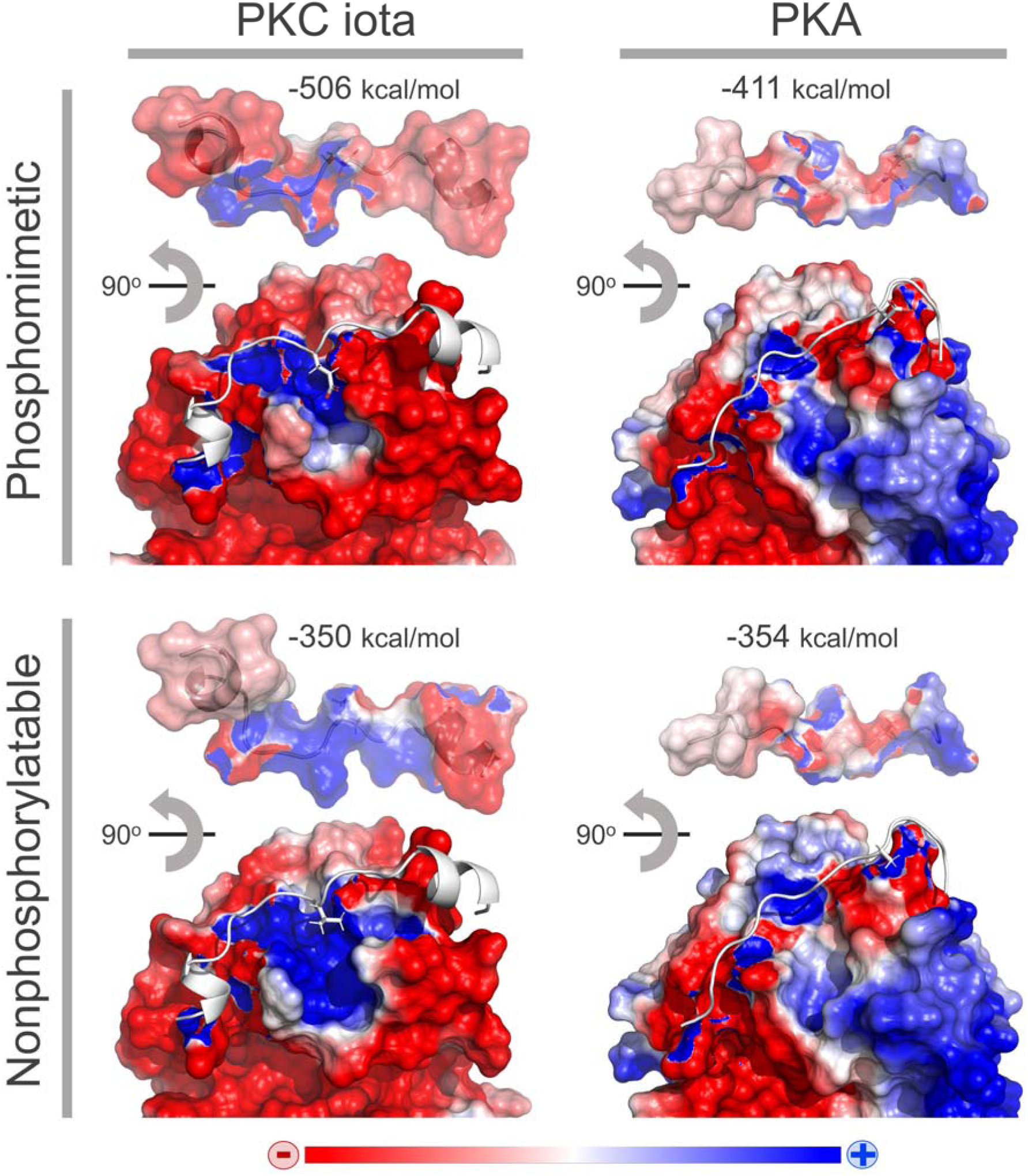
Phosphomimetic (T564E) and nonphosphorylatable (T564V) mutants of PKC iota and phosphomimetic (S339D) and nonphosphorylatable (S339A) mutants of PKA were generated using HADDOCK2.2. For each four mutant, post-simulation structure with the lowest HADDOCK score was subjected to APBS calculations. The surface electrostatics potentials of the proteins were calculated and mapped onto the three-dimensional structures. Red indicates negative potential and blue indicates positive potential. In each structure, C-tail is represented as white cartoon. The 90° rotated state of the C-tails were given on top of their respective structures in order to show the surface electrostatics potentials of the interface. The C-tail binding electrostatics scores that were obtained from HADDOCK2.2 are given on top of each structure. Lower electrostatics score corresponds to stronger binding.

## CONCLUSION

In AGC kinases, the activation loop and C-tail phosphorylations are crucial for kinase activation. Activating phosphorylations govern the conformational transition from inactive to active state through a chain of molecular interactions. A feature that makes Mastl a unique AGC kinase, other than its long NCMR domain, is that, it has an N-lobe hydrophobic groove, yet it is devoid of a hydrophobic motif (HF) (FXXF) to bind it. In other words, Mastl’s C-tail is partially missing. Another protein kinase that shares this unique feature is PDK1. PDK1 circumvents this structural shortfall by binding the HF motifs of other AGC kinases to achieve complete activation (Biondi, 2001; Biondi et al., 2000). Consistently, Vigneron et al. had demonstrated that a partially active mutant form of human Mastl (K72M) could achieve 156% of its basal activity in the presence of a synthetic peptide (PIFtide) that contains hydrophobic motif (Vigneron et al., 2011). Given the lack of HF motif in Mastl and given that Mastl possibly compensates for this by utilizing the HF motif of other kinases, one cannot directly assign a biological function to the C-tail phosphosite.

Although AGC kinases have been studied for decades, role of the C-tail phosphosite in the final activation step is still a mystery. Our experimental results suggest that Mastl pS861 is auxiliary, and not essential for Mastl activation. In line with the experimental findings, our computational results suggest that Mastl pS861 is unlikely to contribute to intramolecular interactions. However, this does not mean that the phosphosite would be redundant, rather it may serve a different function. It is also possible that the phosphosite interacts with a partner protein kinase within a heteromeric complex, which is not proven yet. Aside from our contradicting remarks on the C-tail phosphosite, we support the cis-autophosphorylation model, in which the partially active monomers are responsible for their own hyperphosphorylation.

At this point, it is necessary to experimentally determine the structure of Mastl with its full-length C-tail, and to further explore its possible intramolecular or intermolecular interactions by harnessing structural approaches. More importantly, our present results render the generalizability of the final step of current AGC kinase activation model (Arencibia et al., 2013; Hauge et al., 2007; Kannan et al., 2007) questionable for kinases that are devoid of an HF motif. We hope that our results encourage the researchers in the field to engage in deeper structural and enzymological studies for AGC protein kinases. On the other hand, we have demonstrated the complementation cloning strategy as a robust assay that allows cell culture-based mutagenesis screening of the essential proteins.

## ASSOCIATED CONTENT

### Supporting Information

Supporting information including figures and tables that are mentioned in the text was provided.

## AUTHOR INFORMATION

### Author Contributions

The manuscript was written through contributions of all authors. All authors have given approval to the final version of the manuscript.

### Notes

The authors declare no competing financial interest.

## FUNDING

## Acknowledgements

This work was supported by Dokuz Eylul University Scientific Research Projects grant (DEU-BAP Project No: 2016.KB.SAG.014). Kasim Diril received additional support from Turkish Academy of Sciences (GEBIP award) and The Science Academy, Turkey (BAGEP award). Mehmet Erguven was supported by a stipend from TUBITAK (The Scientific and Technological Research Council of Turkey) Project No. 217Z248

## ACKNOWLEDGEMENTS

We thank Dr. Philipp Kaldis for sharing the plasmids, cell lines and antibodies used in this study.

## ABBREVIATIONS

4-OHT: 4-Hydroxytamoxifen
DMSO: dimethyl sulfoxide
HA: Human influenza hemagglutinin
HF: Hydrophobic Motif
MEF: Mouse mbryonic Fibroblast
NCMR: Non-Conserved Middle Region
NLT: N-Lobe Tether
PDB: The Protein Data Bank
SAC: Spindle Assembly Checkpoint
WT: Wild-Type
KD: Kinase-Dead

## Notes

### Competing Interest Statement

The authors have declared no competing interest.

### Summary of Updates

The units (kcal/mol) for the boxplots (y-axis label) were added in the main manuscript and the supplementary material. Other small/aesthetic changes made in some supplementary figures

