## Supplementary material for "Complementation cloning identifies the essentials of mammalian Mastl kinase activation"

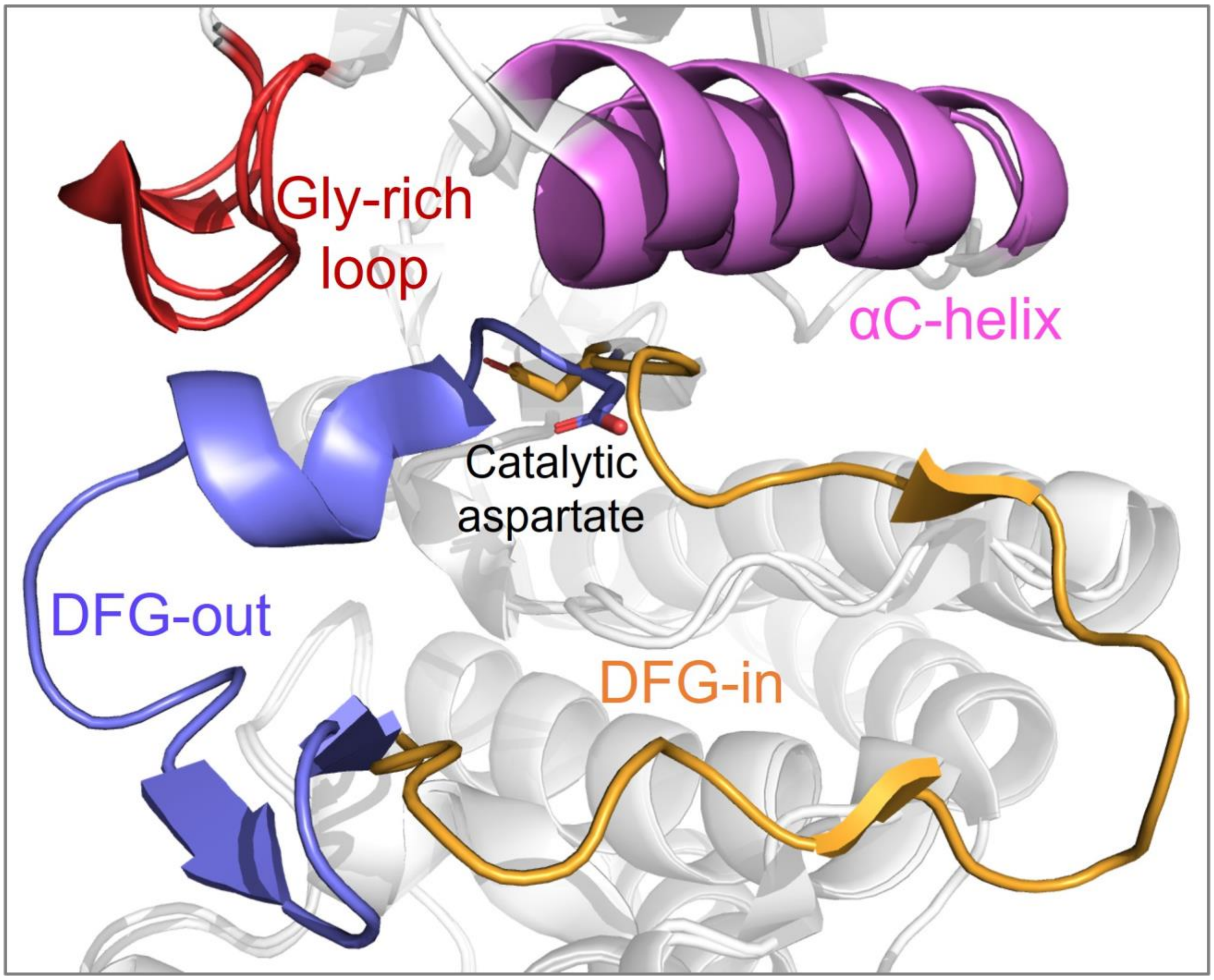


**Figure S1.** The structural organization of protein kinases. The catalytically important segments; glycine-rich loop (red), activation loop (DFG-out is purple, DFG-in is orange), and αC-helix (pink) are represented. DFG-in and DFG-out conformations of Abl kinase were superposed (PDB entries 3KF4 and 3KFA, respectively) (Zhou et al., 2010).

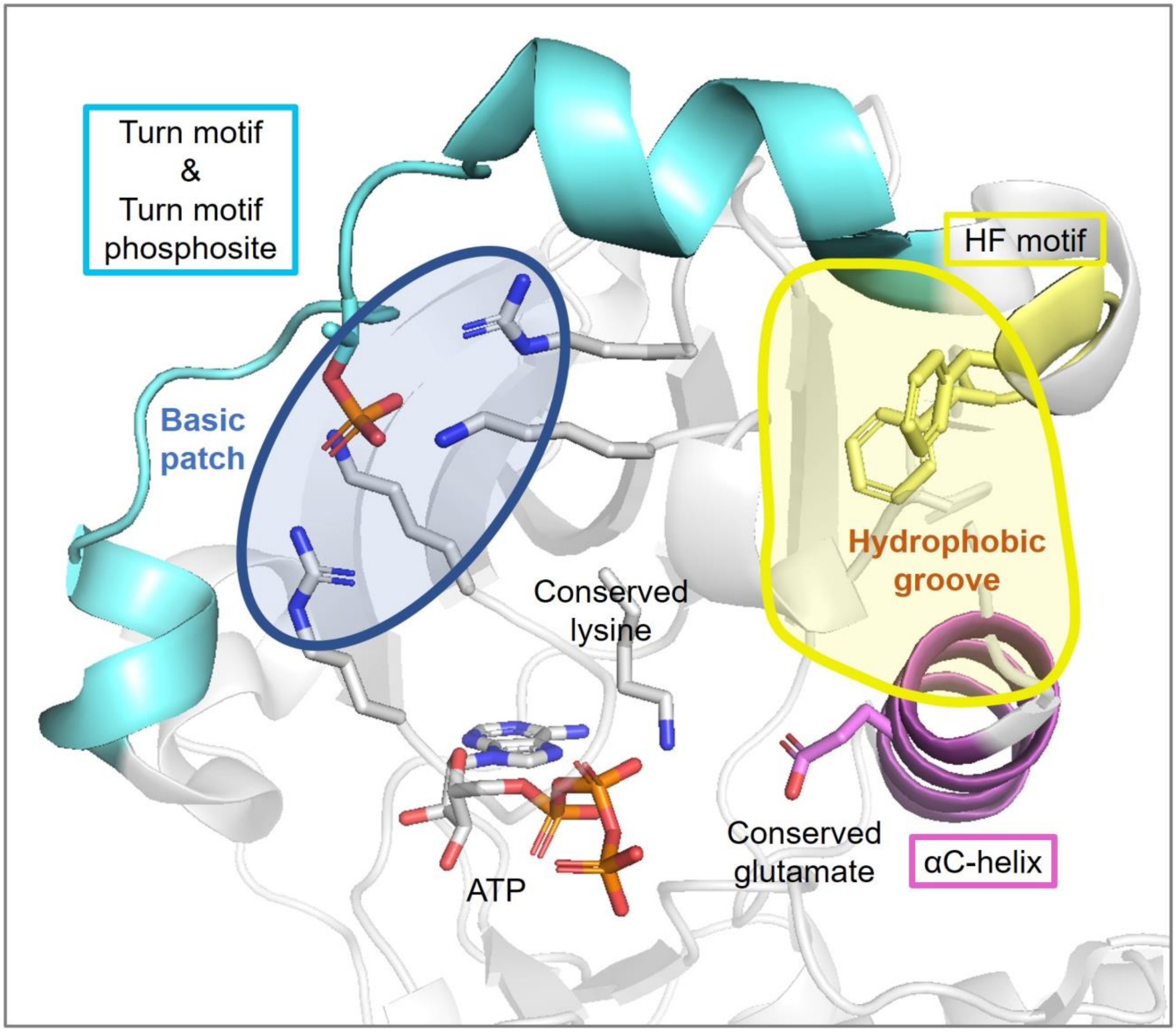


**Figure S2.** The crystal structure of ATP-bound human PKC iota (PDB entry 3A8W) (Takimura et al., 2010). In the N-lobe the phosphoresidue, conserved lysine and glutamate, the HF motif conserved phenylalanine residues, and the residues that form the basic patch are shown in stick forms. The blue, orange, and red colored atoms are nitrogen, phosphorus, and oxygen, respectively. The basic patch and the hydrophobic groove are indicated by the transparent blue and yellow areas, respectively. The turn motif is cyan colored. The αC-helix (magenta), HF motif (yellow), and the proximal β-strand (transparent gray) together form the hydrophobic groove.

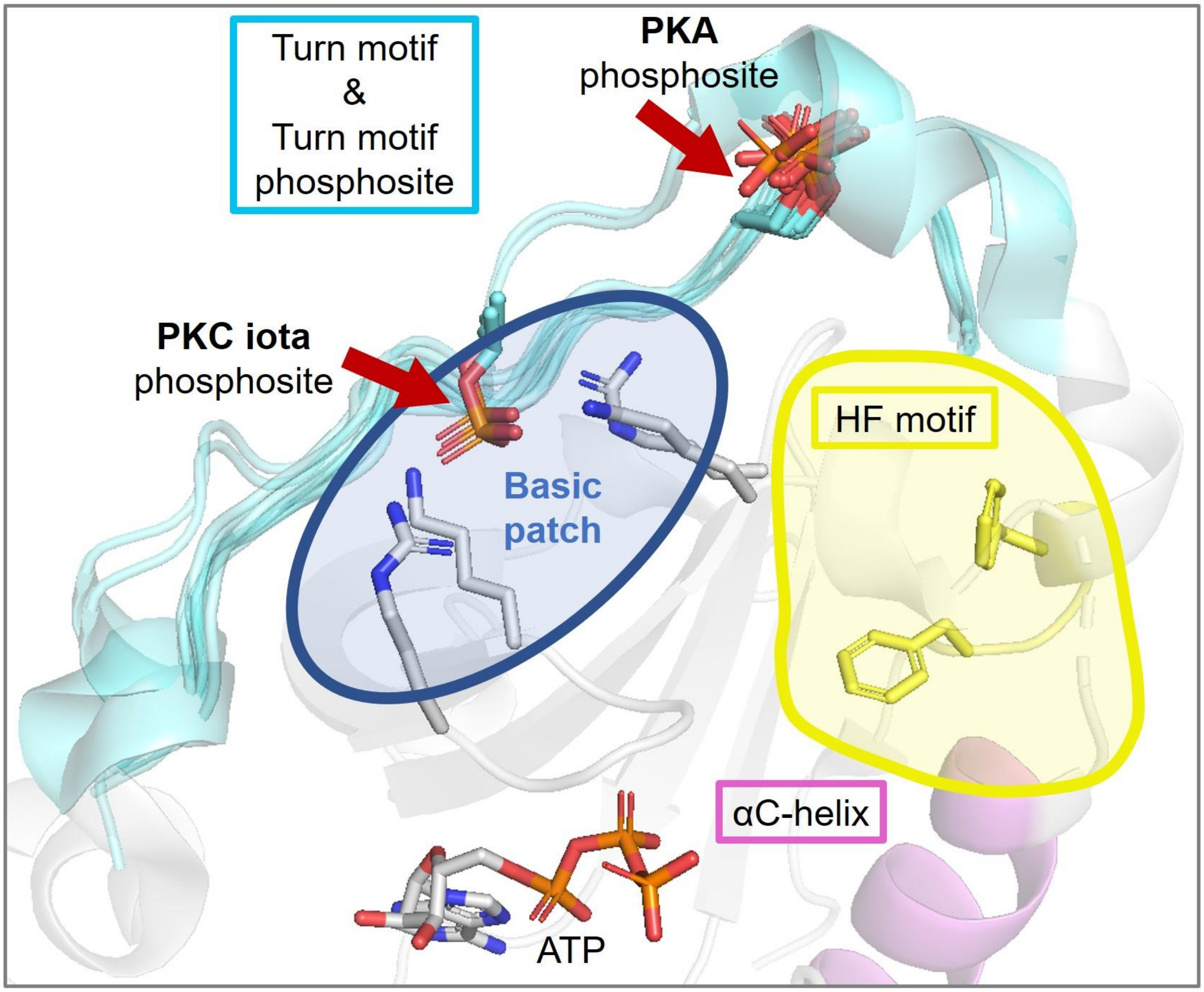


**Figure S3.** A total of 17 AGC kinase structures were obtained from the PDB and superposed in PyMOL. 2 of the structures are PKC iota and 15 of the structures are PKA. The C-tail phosphosite is buried in PKC iota structures while it is solvent exposed in PKA structures. Basic patch is indicated only for one representative PKC iota structure (3A8W). PKA does not possess a corresponding basic patch at its N-lobe surface. The phosphoresidues, basic patch residues, and the HF motif conserved phenylalanine residues are shown in stick forms. The blue, orange, and red colored atoms are nitrogen, phosphorus, and oxygen, respectively. The basic patch and the hydrophobic groove are indicated by the transparent blue and yellow areas, respectively. The turn motif, HF motif, and αC-helix are cyan, yellow, and magenta colored respectively.

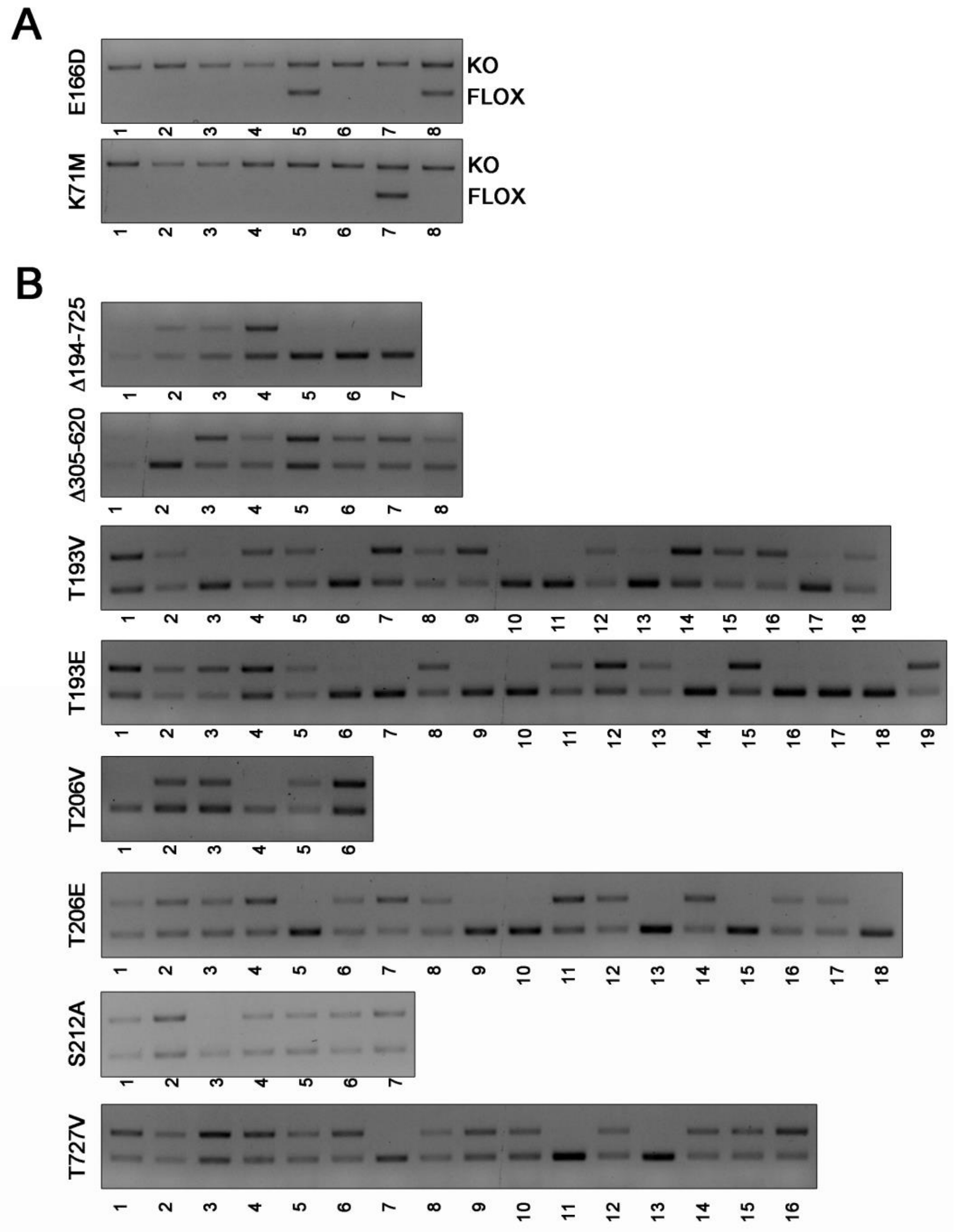


**Figure S4.** Genotype analysis of clonal cell lines expressing different Mastl mutants. After limited dilution, clonal cell lines were generated and analyzed as explained in Figure 2.

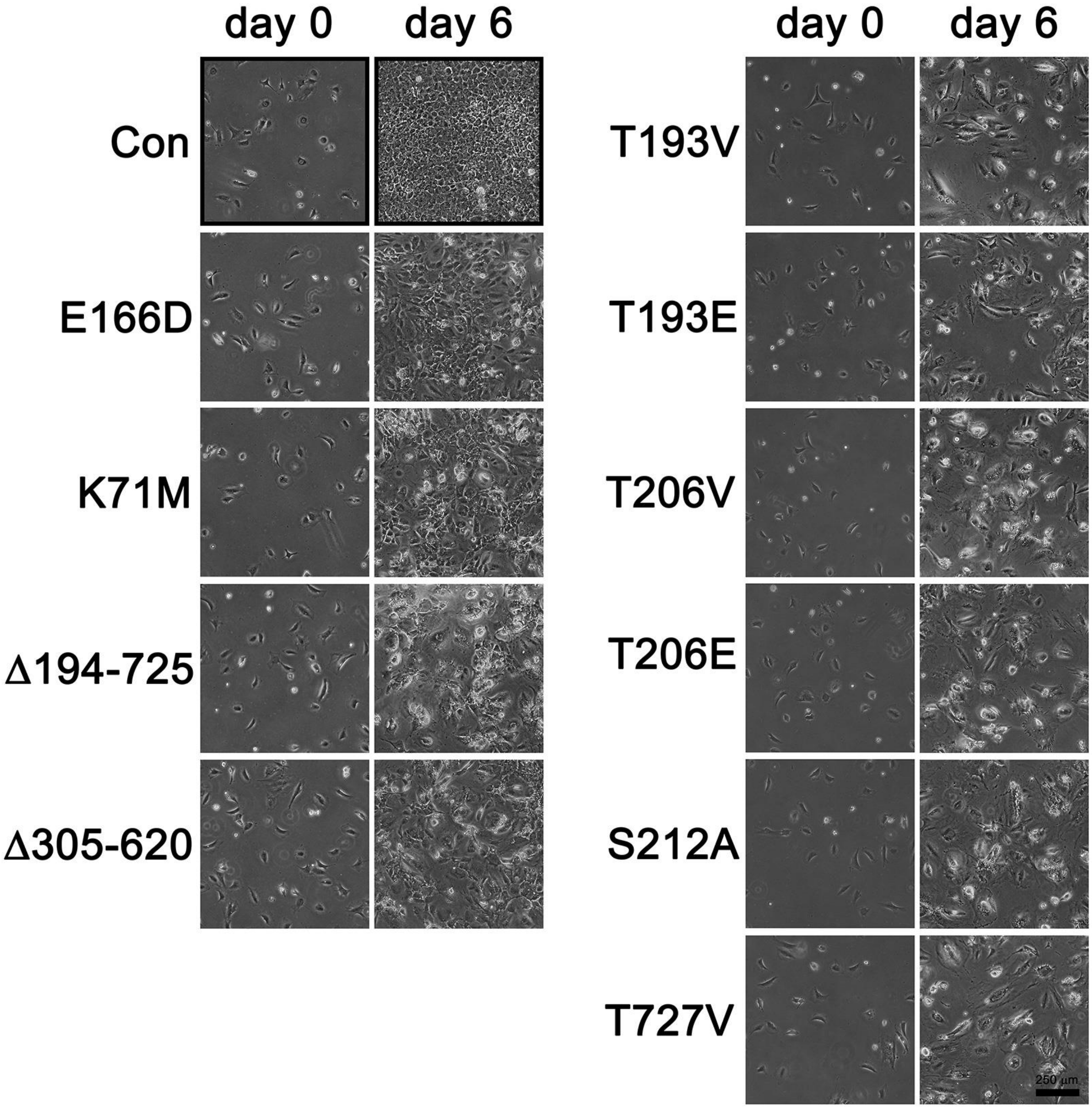


**Figure S5.** Proliferation of stable cell pools after deletion of endogenous Mastl. The micrographs were acquired immediately before (day 0) and six days (day 6) after 4-OHT treatment. Scale bar is 250 µm. Con indicates untreated cells.

```
Mastl      388 NNAQHLTISGFSL------------------     400
               |....||.....:
PKC iota   298 NEPVQLTPDDDDIVRKIDQSEFEGFEYINPL     328

Mastl      381 TSYFEARNNAQHLTISGFSL-----------     400
               ||.|:.... :.:.:   |:
PKA        310 TSNFDDYEE-EEIRV---SINEKCGKEFTEF     336
```

**Figure S6.** Pairwise sequence alignment of the kinase domains of mouse Mastl and human PKC iota or mouse PKA. The alignments were performed by using EMBOSS NEEDLE web server. The NCMR region of Mastl was omitted from the sequence prior to alignment.

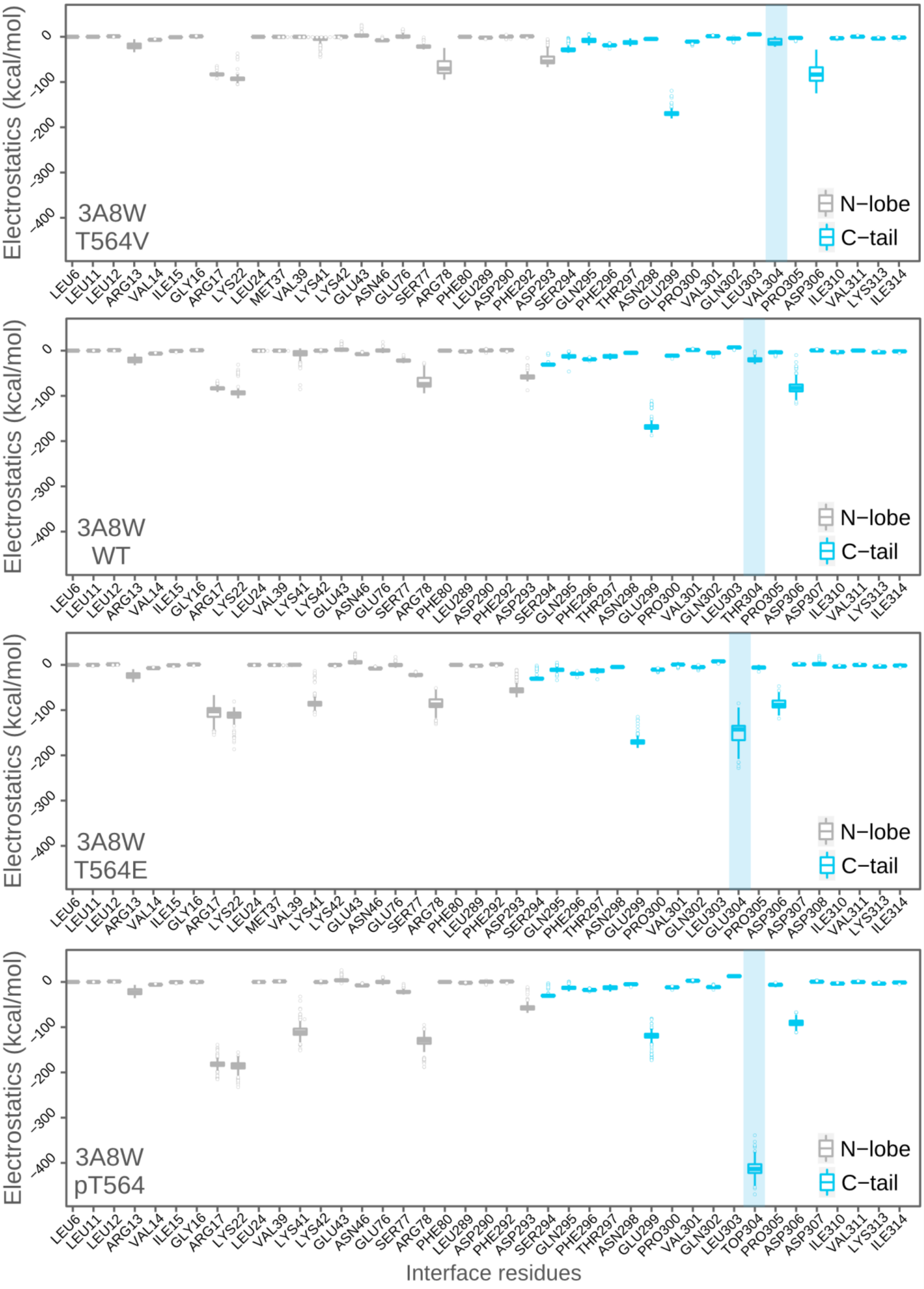

Electrostatics (kcal/mol)
0
-100
-200
-300
-400
3A8W
T564V
N−lobe
C−tail
LEU6 LEU11 LEU12 ARG13 VAL14 ILE15 GLY16 ARG17 LYS22 LEU24 MET37 VAL39 LYS41 LYS42 GLU43 ASN46 GLU76 SER77 ARG78 PHE80 LEU289 ASP290 PHE292 ASP293 SER294 GLN295 PHE296 THR297 ASN298 GLU299 PRO300 VAL301 GLN302 LEU303 VAL304 PRO305 ASP306 ILE310 VAL311 LYS313 ILE314
3A8W
WT
LEU6 LEU11 LEU12 ARG13 VAL14 ILE15 GLY16 ARG17 LYS22 LEU24 VAL39 LYS41 LYS42 GLU43 ASN46 GLU76 SER77 ARG78 PHE80 LEU289 ASP290 PHE292 ASP293 SER294 GLN295 PHE296 THR297 ASN298 GLU299 PRO300 VAL301 GLN302 LEU303 THR304 PRO305 ASP306 ASP307 ILE310 VAL311 LYS313 ILE314
3A8W
T564E
LEU6 LEU11 LEU12 ARG13 VAL14 ILE15 GLY16 ARG17 LYS22 LEU24 MET37 VAL39 LYS41 LYS42 GLU43 ASN46 GLU76 SER77 ARG78 PHE80 LEU289 PHE292 ASP293 SER294 GLN295 PHE296 THR297 ASN298 GLU299 PRO300 VAL301 GLN302 LEU303 GLU304 PRO305 ASP306 ASP307 ASP308 ILE310 VAL311 LYS313 ILE314
3A8W
pT564
LEU6 LEU11 LEU12 ARG13 VAL14 ILE15 GLY16 ARG17 LYS22 LEU24 VAL39 LYS41 LYS42 GLU43 ASN46 GLU76 SER77 ARG78 PHE80 LEU289 ASP290 PHE292 ASP293 SER294 GLN295 PHE296 THR297 ASN298 GLU299 PRO300 VAL301 GLN302 LEU303 TOP304 PRO305 ASP306 ASP307 ILE310 VAL311 LYS313 ILE314
Interface residues

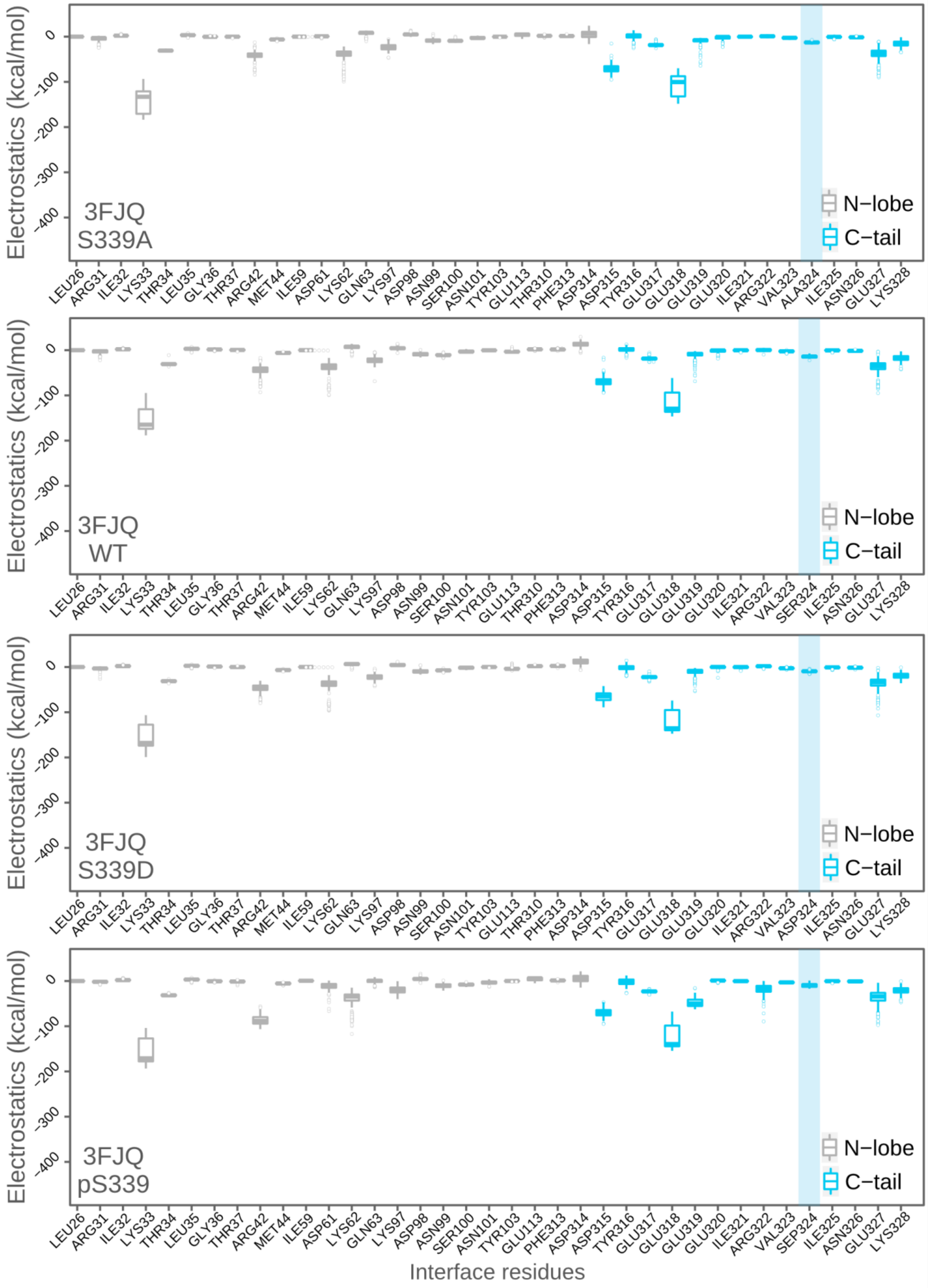
3FJQ S339A
3FJQ WT
3FJQ S339D
3FJQ pS339
N−lobe
C−tail
Electrostatics (kcal/mol)
0
-100
-200
-300
-400
LEU26 ARG31 ILE32 LYS33 THR34 LEU35 GLY36 THR37 ARG42 MET44 ILE59 ASP61 LYS62 GLN63 LYS97 ASP98 ASN99 SER100 ASN101 TYR103 GLU113 THR310 PHE313 ASP314 ASP315 TYR316 GLU317 GLU318 GLU319 GLU320 ILE321 ARG322 VAL323 ALA324 ILE325 ASN326 GLU327 LYS328
SER324
ASP324
SEP324
Interface residues

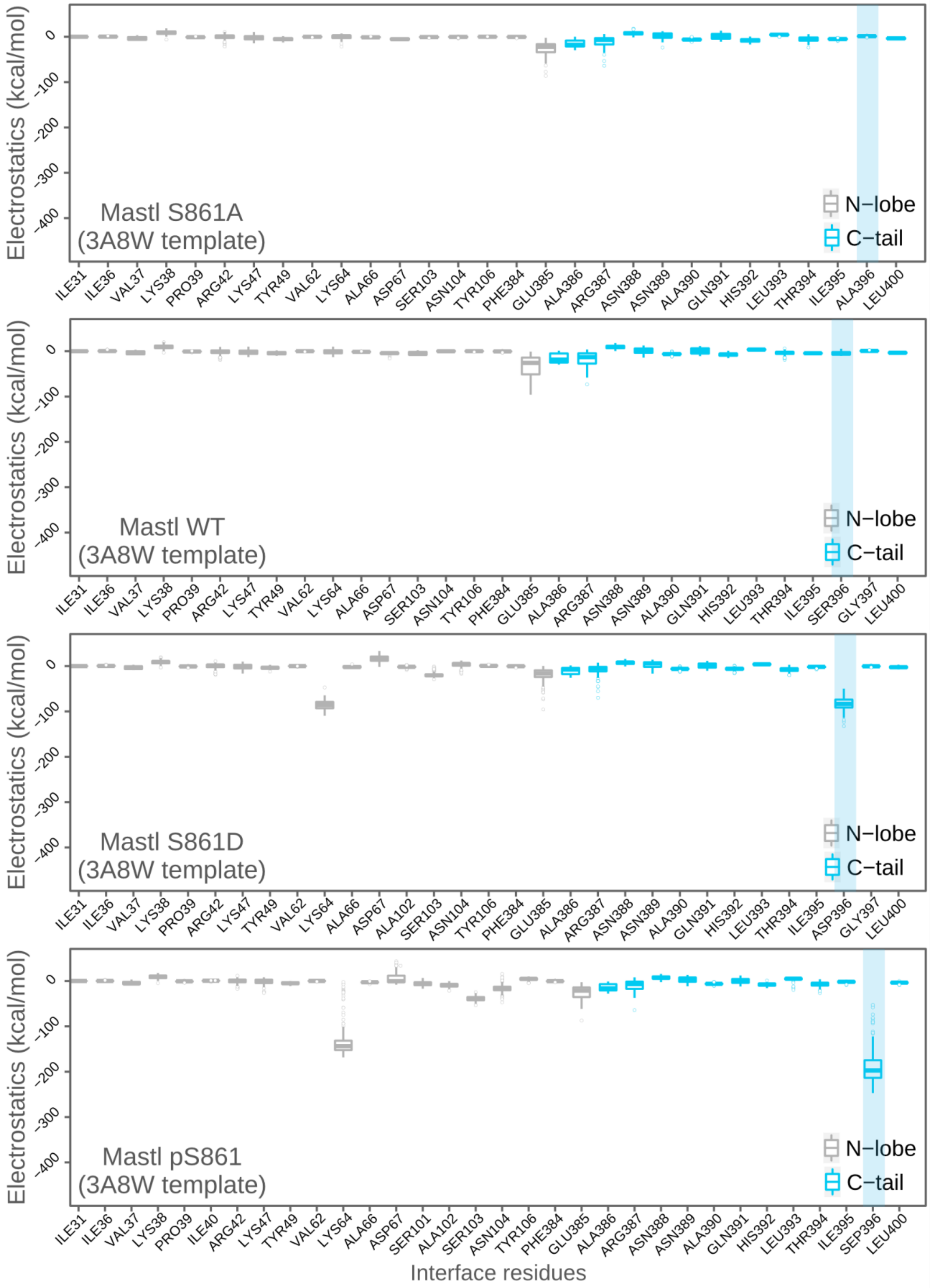

Electrostatics (kcal/mol)
Mastl S861A
(3A8W template)
N−lobe
C−tail
ILE31 ILE36 VAL37 LYS38 PRO39 ARG42 LYS47 TYR49 VAL62 LYS64 ALA66 ASP67 SER103 ASN104 TYR106 PHE384 GLU385 ALA386 ARG387 ASN388 ASN389 ALA390 GLN391 HIS392 LEU393 THR394 ILE395 ALA396 LEU400
Mastl WT
(3A8W template)
ILE31 ILE36 VAL37 LYS38 PRO39 ARG42 LYS47 TYR49 VAL62 LYS64 ALA66 ASP67 SER103 ASN104 TYR106 PHE384 GLU385 ALA386 ARG387 ASN388 ASN389 ALA390 GLN391 HIS392 LEU393 THR394 ILE395 SER396 GLY397 LEU400
Mastl S861D
(3A8W template)
ILE31 ILE36 VAL37 LYS38 PRO39 ARG42 LYS47 TYR49 VAL62 LYS64 ALA66 ASP67 ALA102 SER103 ASN104 TYR106 PHE384 GLU385 ALA386 ARG387 ASN388 ASN389 ALA390 GLN391 HIS392 LEU393 THR394 ILE395 ASP396 GLY397 LEU400
Mastl pS861
(3A8W template)
ILE31 ILE36 VAL37 LYS38 PRO39 ILE40 ARG42 LYS47 TYR49 VAL62 LYS64 ALA66 ASP67 SER101 ALA102 SER103 ASN104 TYR106 PHE384 GLU385 ALA386 ARG387 ASN388 ASN389 ALA390 GLN391 HIS392 LEU393 THR394 ILE395 SEP396 LEU400
Interface residues
0
-100
-200
-300
-400

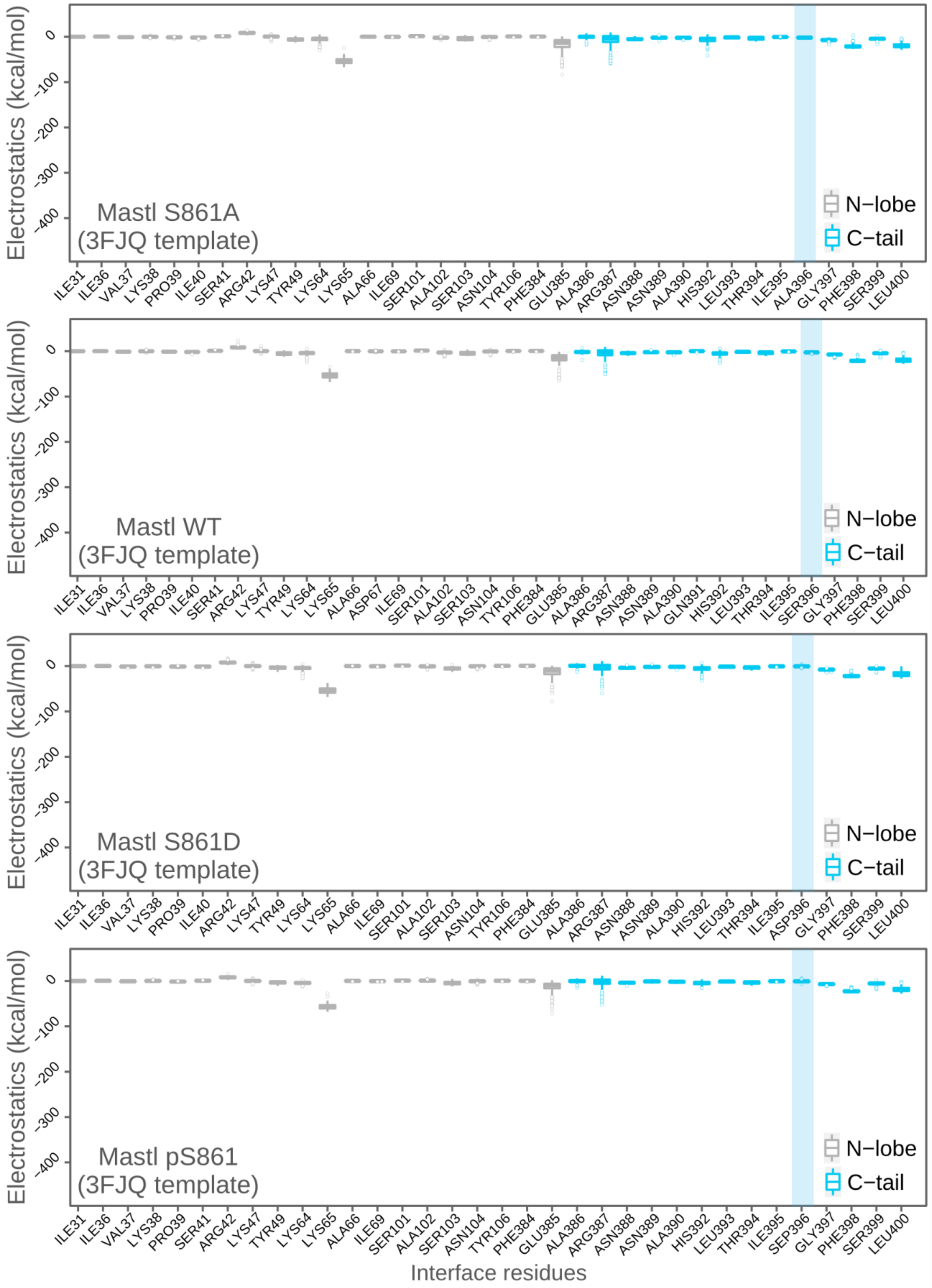
Electrostatics (kcal/mol)
0
-100
-200
-300
-400
Mastl S861A
(3FJQ template)
N−lobe
C−tail
ILE31 ILE36 VAL37 LYS38 PRO39 ILE40 SER41 ARG42 LYS47 TYR49 LYS64 LYS65 ALA66 ILE69 SER101 ALA102 SER103 ASN104 TYR106 PHE384 GLU385 ALA386 ARG387 ASN388 ASN389 ALA390 HIS392 LEU393 THR394 ILE395 ALA396 GLY397 PHE398 SER399 LEU400
Electrostatics (kcal/mol)
0
-100
-200
-300
-400
Mastl WT
(3FJQ template)
N−lobe
C−tail
ILE31 ILE36 VAL37 LYS38 PRO39 ILE40 SER41 ARG42 LYS47 TYR49 LYS64 LYS65 ALA66 ASP67 ILE69 SER101 ALA102 SER103 ASN104 TYR106 PHE384 GLU385 ALA386 ARG387 ASN388 ASN389 ALA390 GLN391 HIS392 LEU393 THR394 ILE395 SER396 GLY397 PHE398 SER399 LEU400
Electrostatics (kcal/mol)
0
-100
-200
-300
-400
Mastl S861D
(3FJQ template)
N−lobe
C−tail
ILE31 ILE36 VAL37 LYS38 PRO39 ILE40 ARG42 LYS47 TYR49 LYS64 LYS65 ALA66 ILE69 SER101 ALA102 SER103 ASN104 TYR106 PHE384 GLU385 ALA386 ARG387 ASN388 ASN389 ALA390 HIS392 LEU393 THR394 ILE395 ASP396 GLY397 PHE398 SER399 LEU400
Electrostatics (kcal/mol)
0
-100
-200
-300
-400
Mastl pS861
(3FJQ template)
N−lobe
C−tail
ILE31 ILE36 VAL37 LYS38 PRO39 SER41 ARG42 LYS47 TYR49 LYS64 LYS65 ALA66 ILE69 SER101 ALA102 SER103 ASN104 TYR106 PHE384 GLU385 ALA386 ARG387 ASN388 ASN389 ALA390 HIS392 LEU393 THR394 ILE395 SEP396 GLY397 PHE398 SER399 LEU400
Interface residues

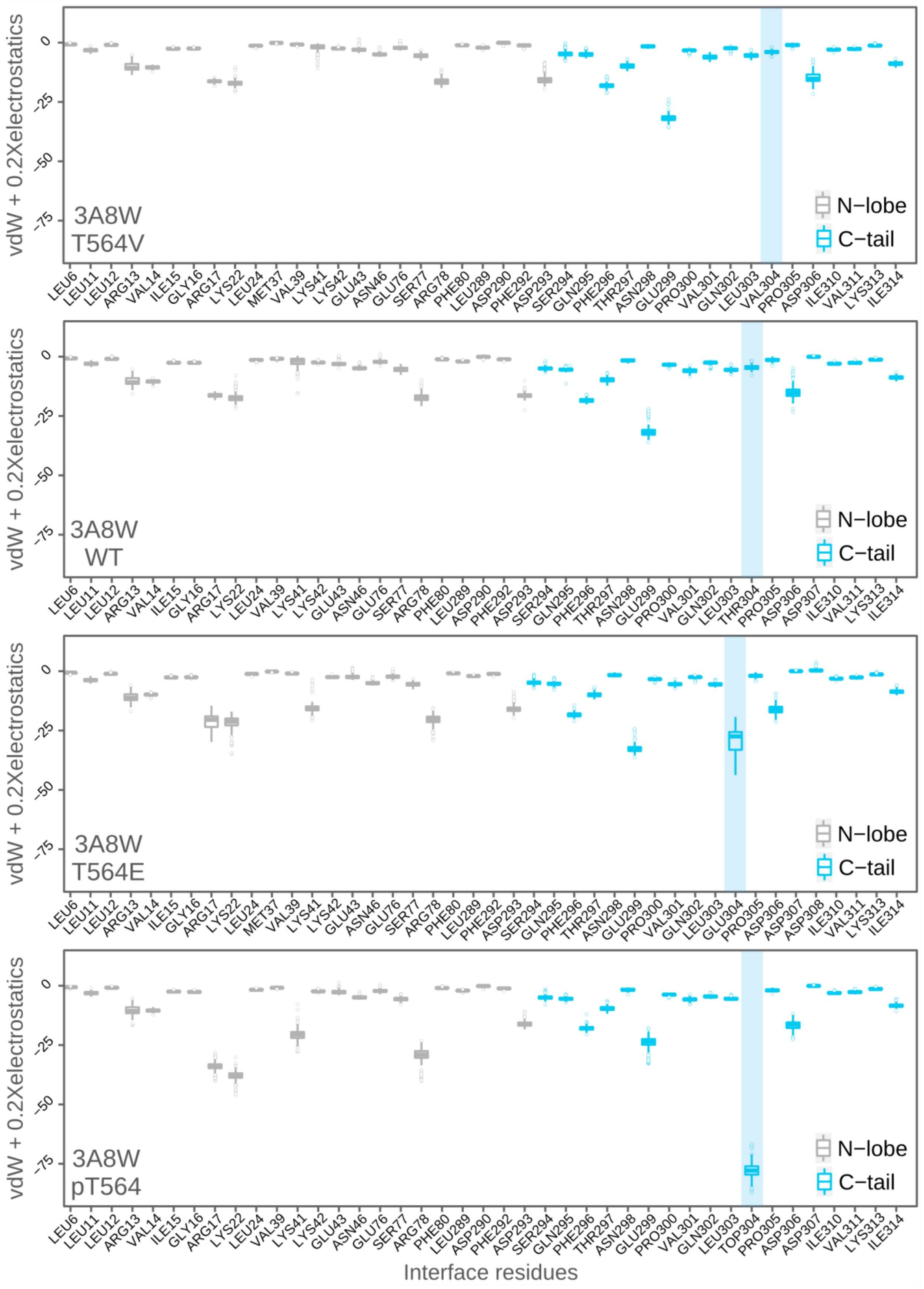
vdW + 0.2Xelectrostatics
0
-25
-50
-75
3A8W T564V
N−lobe
C−tail
LEU6 LEU11 LEU12 ARG13 VAL14 ILE15 GLY16 ARG17 LYS22 LEU24 MET37 VAL39 LYS41 LYS42 GLU43 ASN46 GLU76 SER77 ARG78 PHE80 LEU289 ASP290 PHE292 ASP293 SER294 GLN295 PHE296 THR297 ASN298 GLU299 PRO300 VAL301 GLN302 LEU303 VAL304 PRO305 ASP306 ILE310 VAL311 LYS313 ILE314
3A8W WT
LEU6 LEU11 LEU12 ARG13 VAL14 ILE15 GLY16 ARG17 LYS22 LEU24 VAL39 LYS41 LYS42 GLU43 ASN46 GLU76 SER77 ARG78 PHE80 LEU289 ASP290 PHE292 ASP293 SER294 GLN295 PHE296 THR297 ASN298 GLU299 PRO300 VAL301 GLN302 LEU303 THR304 PRO305 ASP306 ASP307 ILE310 VAL311 LYS313 ILE314
3A8W T564E
LEU6 LEU11 LEU12 ARG13 VAL14 ILE15 GLY16 ARG17 LYS22 LEU24 MET37 VAL39 LYS41 LYS42 GLU43 ASN46 GLU76 SER77 ARG78 PHE80 LEU289 PHE292 ASP293 SER294 GLN295 PHE296 THR297 ASN298 GLU299 PRO300 VAL301 GLN302 LEU303 GLU304 PRO305 ASP306 ASP307 ASP308 ILE310 VAL311 LYS313 ILE314
3A8W pT564
LEU6 LEU11 LEU12 ARG13 VAL14 ILE15 GLY16 ARG17 LYS22 LEU24 VAL39 LYS41 LYS42 GLU43 ASN46 GLU76 SER77 ARG78 PHE80 LEU289 ASP290 PHE292 ASP293 SER294 GLN295 PHE296 THR297 ASN298 GLU299 PRO300 VAL301 GLN302 LEU303 TOP304 PRO305 ASP306 ASP307 ILE310 VAL311 LYS313 ILE314
Interface residues

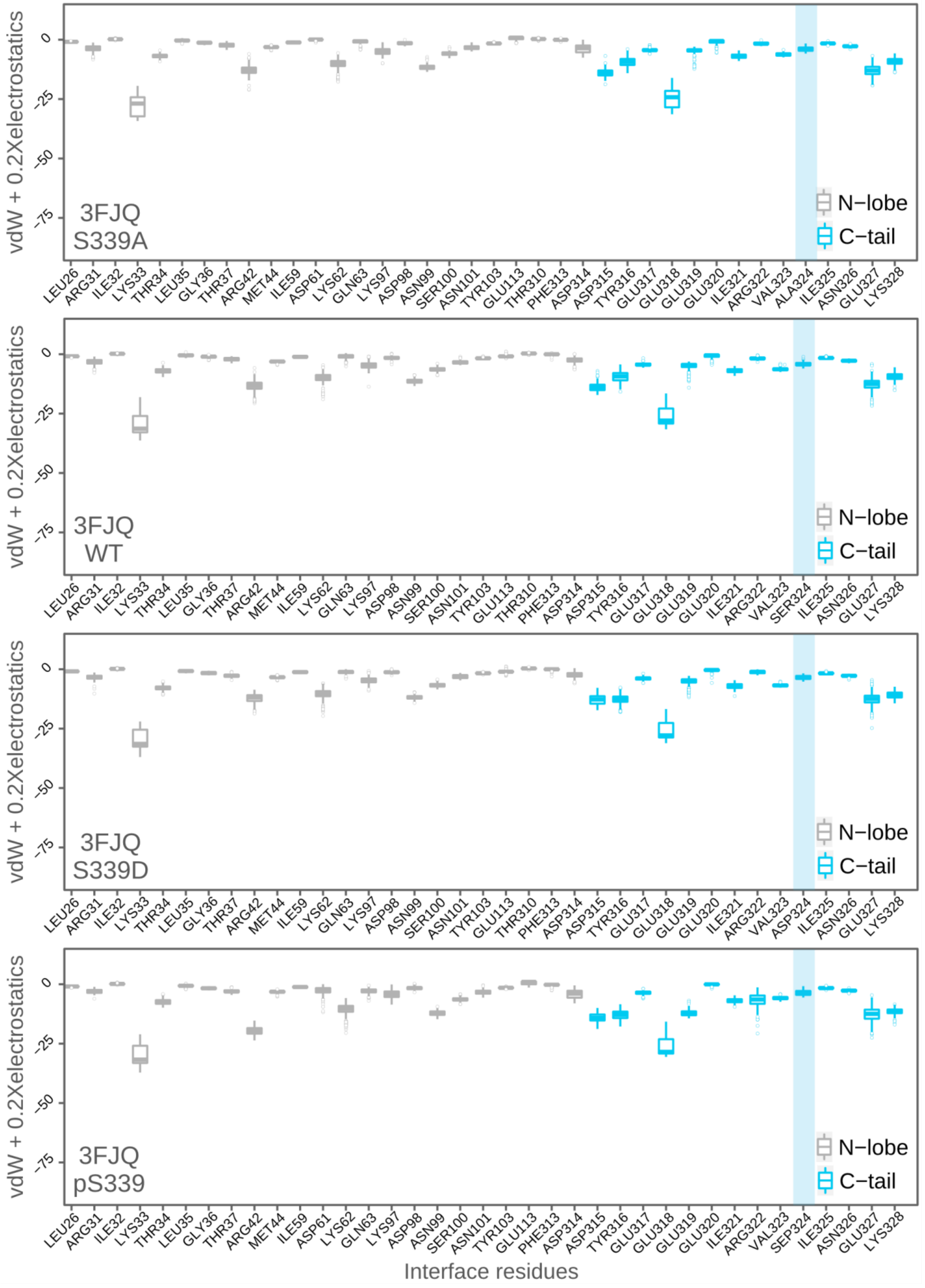
vdW + 0.2Xelectrostatics
0
-25
-50
-75
3FJQ
S339A
N−lobe
C−tail
LEU26 ARG31 ILE32 LYS33 THR34 LEU35 GLY36 THR37 ARG42 MET44 ILE59 ASP61 LYS62 GLN63 LYS97 ASP98 ASN99 SER100 ASN101 TYR103 GLU113 THR310 PHE313 ASP314 ASP315 TYR316 GLU317 GLU318 GLU319 GLU320 ILE321 ARG322 VAL323 ALA324 ILE325 ASN326 GLU327 LYS328
3FJQ
WT
SER324
3FJQ
S339D
ASP324
3FJQ
pS339
SEP324
Interface residues

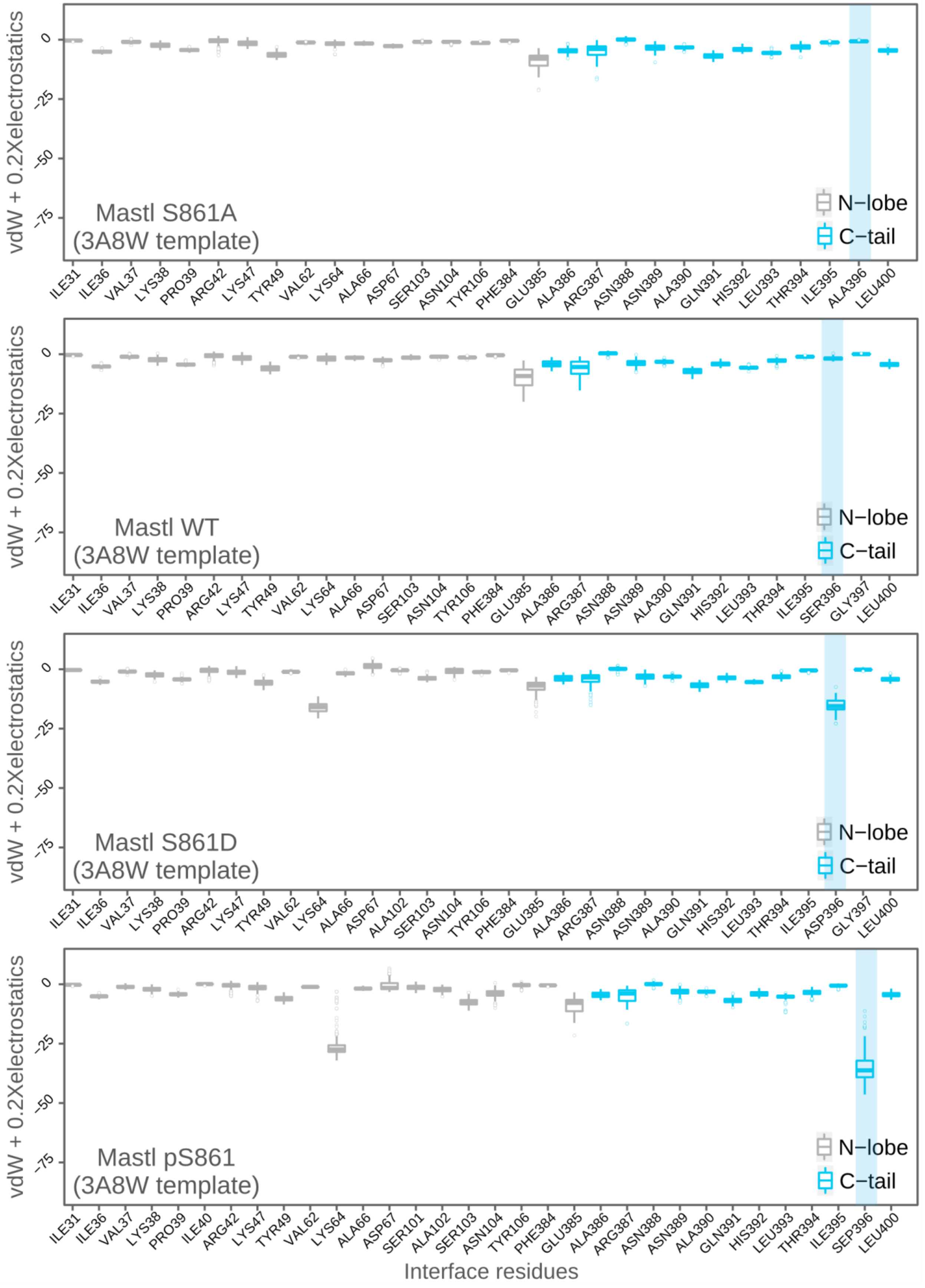
vdW + 0.2Xelectrostatics
0
-25
-50
-75
Mastl S861A
(3A8W template)
N−lobe
C−tail
ILE31 ILE36 VAL37 LYS38 PRO39 ARG42 LYS47 TYR49 VAL62 LYS64 ALA66 ASP67 SER103 ASN104 TYR106 PHE384 GLU385 ALA386 ARG387 ASN388 ASN389 ALA390 GLN391 HIS392 LEU393 THR394 ILE395 ALA396 LEU400
Mastl WT
(3A8W template)
ILE31 ILE36 VAL37 LYS38 PRO39 ARG42 LYS47 TYR49 VAL62 LYS64 ALA66 ASP67 SER103 ASN104 TYR106 PHE384 GLU385 ALA386 ARG387 ASN388 ASN389 ALA390 GLN391 HIS392 LEU393 THR394 ILE395 SER396 GLY397 LEU400
Mastl S861D
(3A8W template)
ILE31 ILE36 VAL37 LYS38 PRO39 ARG42 LYS47 TYR49 VAL62 LYS64 ALA66 ASP67 ALA102 SER103 ASN104 TYR106 PHE384 GLU385 ALA386 ARG387 ASN388 ASN389 ALA390 GLN391 HIS392 LEU393 THR394 ILE395 ASP396 GLY397 LEU400
Mastl pS861
(3A8W template)
ILE31 ILE36 VAL37 LYS38 PRO39 ILE40 ARG42 LYS47 TYR49 VAL62 LYS64 ALA66 ASP67 SER101 ALA102 SER103 ASN104 TYR106 PHE384 GLU385 ALA386 ARG387 ASN388 ASN389 ALA390 GLN391 HIS392 LEU393 THR394 ILE395 SEP396 LEU400
Interface residues

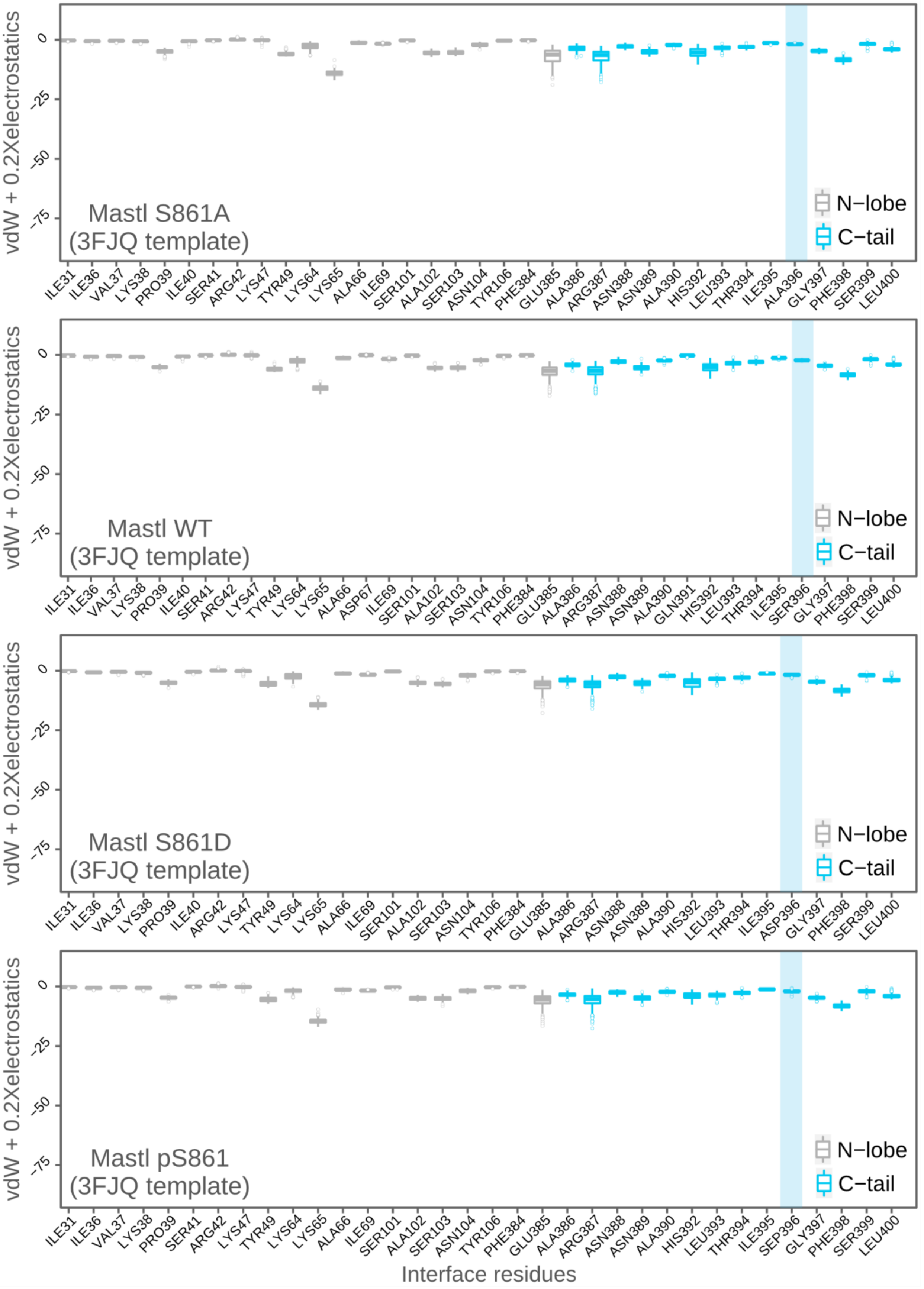
vdW + 0.2Xelectrostatics
0
-25
-50
-75
Mastl S861A
(3FJQ template)
N−lobe
C−tail
ILE31 ILE36 VAL37 LYS38 PRO39 ILE40 SER41 ARG42 LYS47 TYR49 LYS64 LYS65 ALA66 ILE69 SER101 ALA102 SER103 ASN104 TYR106 PHE384 GLU385 ALA386 ARG387 ASN388 ASN389 ALA390 HIS392 LEU393 THR394 ILE395 ALA396 GLY397 PHE398 SER399 LEU400
vdW + 0.2Xelectrostatics
0
-25
-50
-75
Mastl WT
(3FJQ template)
N−lobe
C−tail
ILE31 ILE36 VAL37 LYS38 PRO39 ILE40 SER41 ARG42 LYS47 TYR49 LYS64 LYS65 ALA66 ASP67 ILE69 SER101 ALA102 SER103 ASN104 TYR106 PHE384 GLU385 ALA386 ARG387 ASN388 ASN389 ALA390 GLN391 HIS392 LEU393 THR394 ILE395 SER396 GLY397 PHE398 SER399 LEU400
vdW + 0.2Xelectrostatics
0
-25
-50
-75
Mastl S861D
(3FJQ template)
N−lobe
C−tail
ILE31 ILE36 VAL37 LYS38 PRO39 ILE40 ARG42 LYS47 TYR49 LYS64 LYS65 ALA66 ILE69 SER101 ALA102 SER103 ASN104 TYR106 PHE384 GLU385 ALA386 ARG387 ASN388 ASN389 ALA390 HIS392 LEU393 THR394 ILE395 ASP396 GLY397 PHE398 SER399 LEU400
vdW + 0.2Xelectrostatics
0
-25
-50
-75
Mastl pS861
(3FJQ template)
N−lobe
C−tail
ILE31 ILE36 VAL37 LYS38 PRO39 SER41 ARG42 LYS47 TYR49 LYS64 LYS65 ALA66 ILE69 SER101 ALA102 SER103 ASN104 TYR106 PHE384 GLU385 ALA386 ARG387 ASN388 ASN389 ALA390 HIS392 LEU393 THR394 ILE395 SEP396 GLY397 PHE398 SER399 LEU400
Interface residues

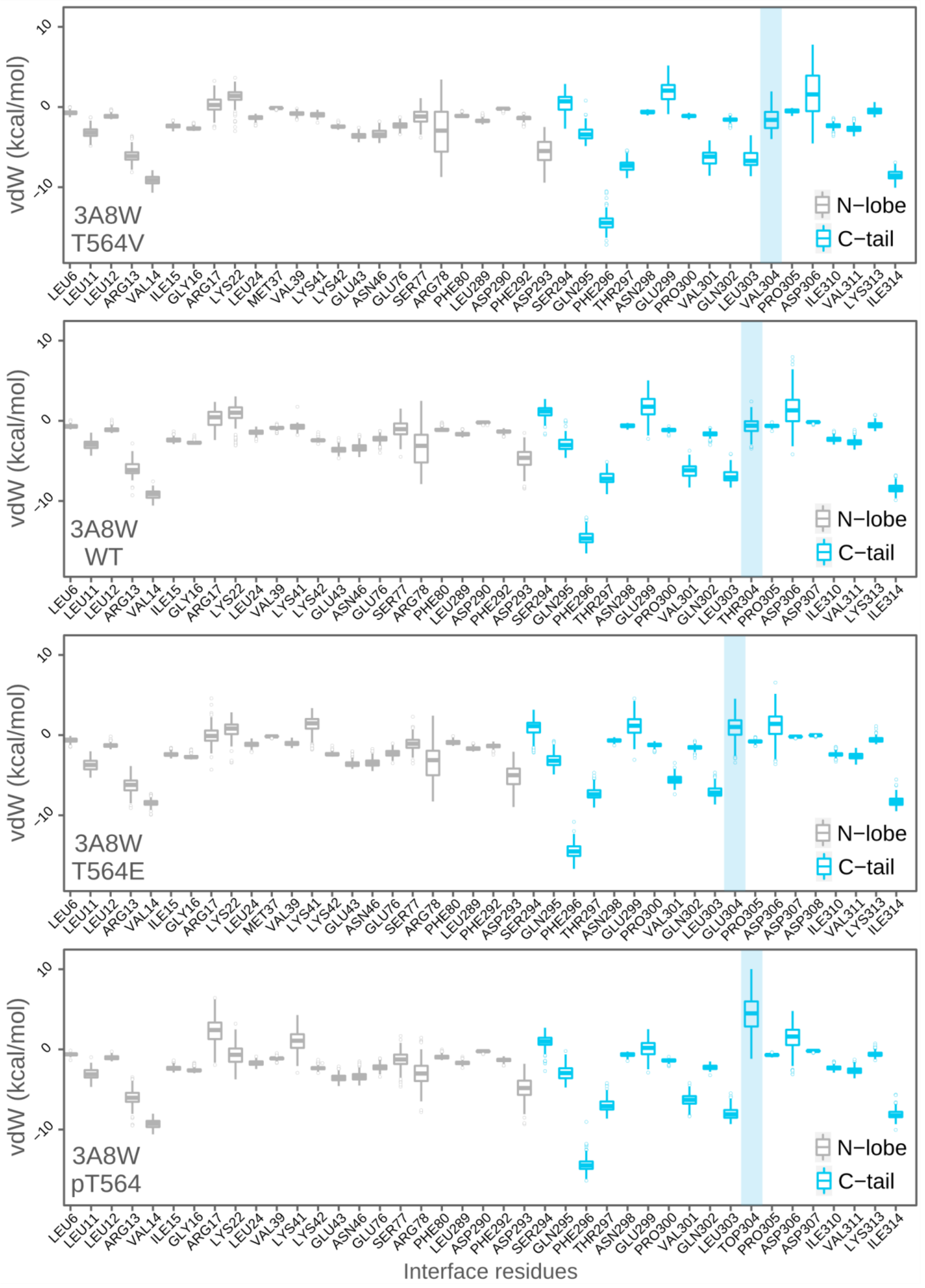
vdW (kcal/mol)
3A8W
T564V
N−lobe
C−tail
LEU6 LEU11 LEU12 ARG13 VAL14 ILE15 GLY16 ARG17 LYS22 LEU24 MET37 VAL39 LYS41 LYS42 GLU43 ASN46 GLU76 SER77 ARG78 PHE80 LEU289 ASP290 PHE292 ASP293 SER294 GLN295 PHE296 THR297 ASN298 GLU299 PRO300 VAL301 GLN302 LEU303 VAL304 PRO305 ASP306 ILE310 VAL311 LYS313 ILE314
3A8W
WT
LEU6 LEU11 LEU12 ARG13 VAL14 ILE15 GLY16 ARG17 LYS22 LEU24 VAL39 LYS41 LYS42 GLU43 ASN46 GLU76 SER77 ARG78 PHE80 LEU289 ASP290 PHE292 ASP293 SER294 GLN295 PHE296 THR297 ASN298 GLU299 PRO300 VAL301 GLN302 LEU303 THR304 PRO305 ASP306 ASP307 ILE310 VAL311 LYS313 ILE314
3A8W
T564E
LEU6 LEU11 LEU12 ARG13 VAL14 ILE15 GLY16 ARG17 LYS22 LEU24 MET37 VAL39 LYS41 LYS42 GLU43 ASN46 GLU76 SER77 ARG78 PHE80 LEU289 PHE292 ASP293 SER294 GLN295 PHE296 THR297 ASN298 GLU299 PRO300 VAL301 GLN302 LEU303 GLU304 PRO305 ASP306 ASP307 ASP308 ILE310 VAL311 LYS313 ILE314
3A8W
pT564
LEU6 LEU11 LEU12 ARG13 VAL14 ILE15 GLY16 ARG17 LYS22 LEU24 VAL39 LYS41 LYS42 GLU43 ASN46 GLU76 SER77 ARG78 PHE80 LEU289 ASP290 PHE292 ASP293 SER294 GLN295 PHE296 THR297 ASN298 GLU299 PRO300 VAL301 GLN302 LEU303 TOP304 PRO305 ASP306 ASP307 ILE310 VAL311 LYS313 ILE314
Interface residues

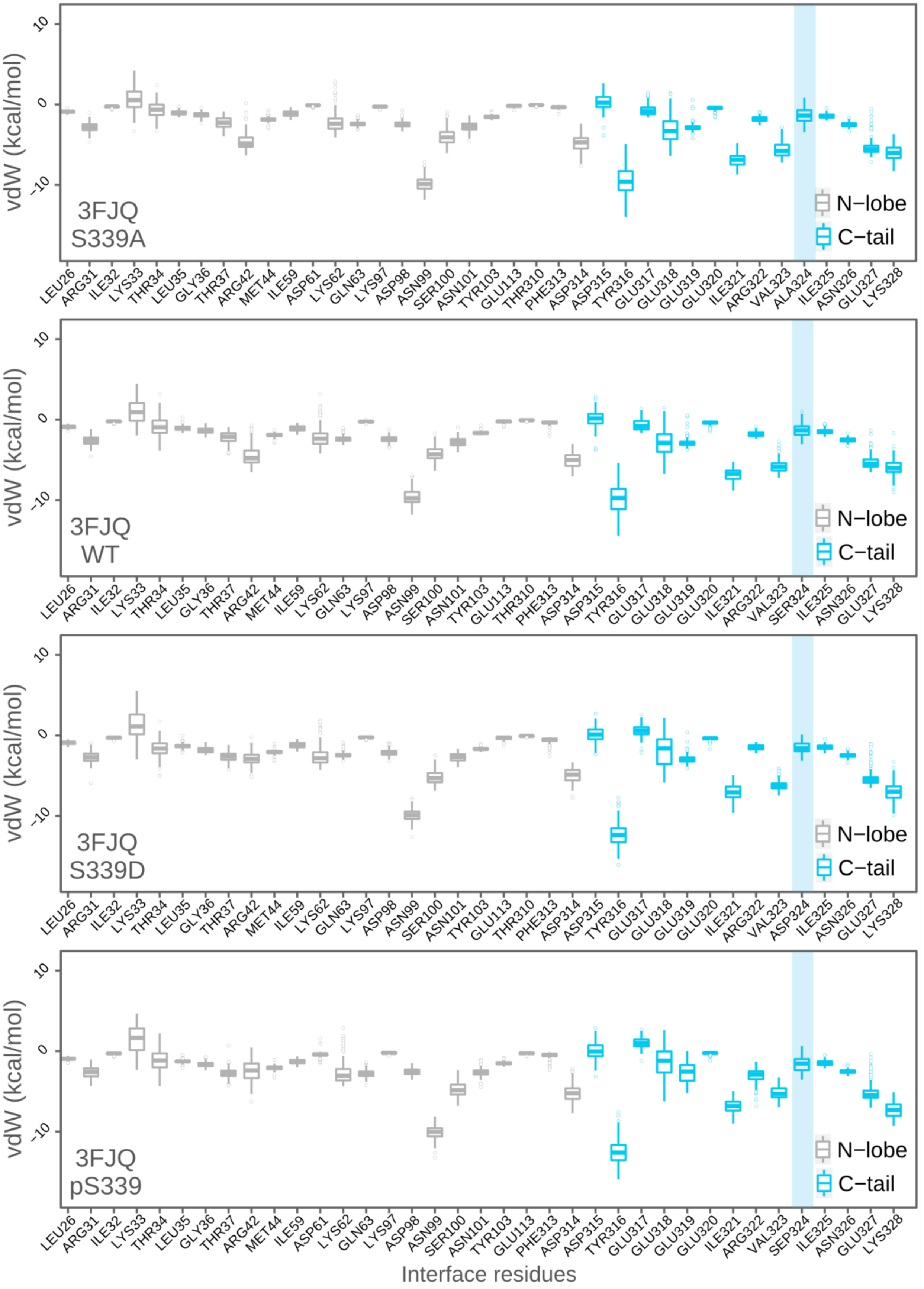

vdW (kcal/mol)
10
0
-10
3FJQ
S339A
N−lobe
C−tail
LEU26 ARG31 ILE32 LYS33 THR34 LEU35 GLY36 THR37 ARG42 MET44 ILE59 ASP61 LYS62 GLN63 LYS97 ASP98 ASN99 SER100 ASN101 TYR103 GLU113 THR310 PHE313 ASP314 ASP315 TYR316 GLU317 GLU318 GLU319 GLU320 ILE321 ARG322 VAL323 ALA324 ILE325 ASN326 GLU327 LYS328
3FJQ
WT
LEU26 ARG31 ILE32 LYS33 THR34 LEU35 GLY36 THR37 ARG42 MET44 ILE59 LYS62 GLN63 LYS97 ASP98 ASN99 SER100 ASN101 TYR103 GLU113 THR310 PHE313 ASP314 ASP315 TYR316 GLU317 GLU318 GLU319 GLU320 ILE321 ARG322 VAL323 SER324 ILE325 ASN326 GLU327 LYS328
3FJQ
S339D
LEU26 ARG31 ILE32 LYS33 THR34 LEU35 GLY36 THR37 ARG42 MET44 ILE59 LYS62 GLN63 LYS97 ASP98 ASN99 SER100 ASN101 TYR103 GLU113 THR310 PHE313 ASP314 ASP315 TYR316 GLU317 GLU318 GLU319 GLU320 ILE321 ARG322 VAL323 ASP324 ILE325 ASN326 GLU327 LYS328
3FJQ
pS339
LEU26 ARG31 ILE32 LYS33 THR34 LEU35 GLY36 THR37 ARG42 MET44 ILE59 ASP61 LYS62 GLN63 LYS97 ASP98 ASN99 SER100 ASN101 TYR103 GLU113 PHE313 ASP314 ASP315 TYR316 GLU317 GLU318 GLU319 GLU320 ILE321 ARG322 VAL323 SEP324 ILE325 ASN326 GLU327 LYS328
Interface residues

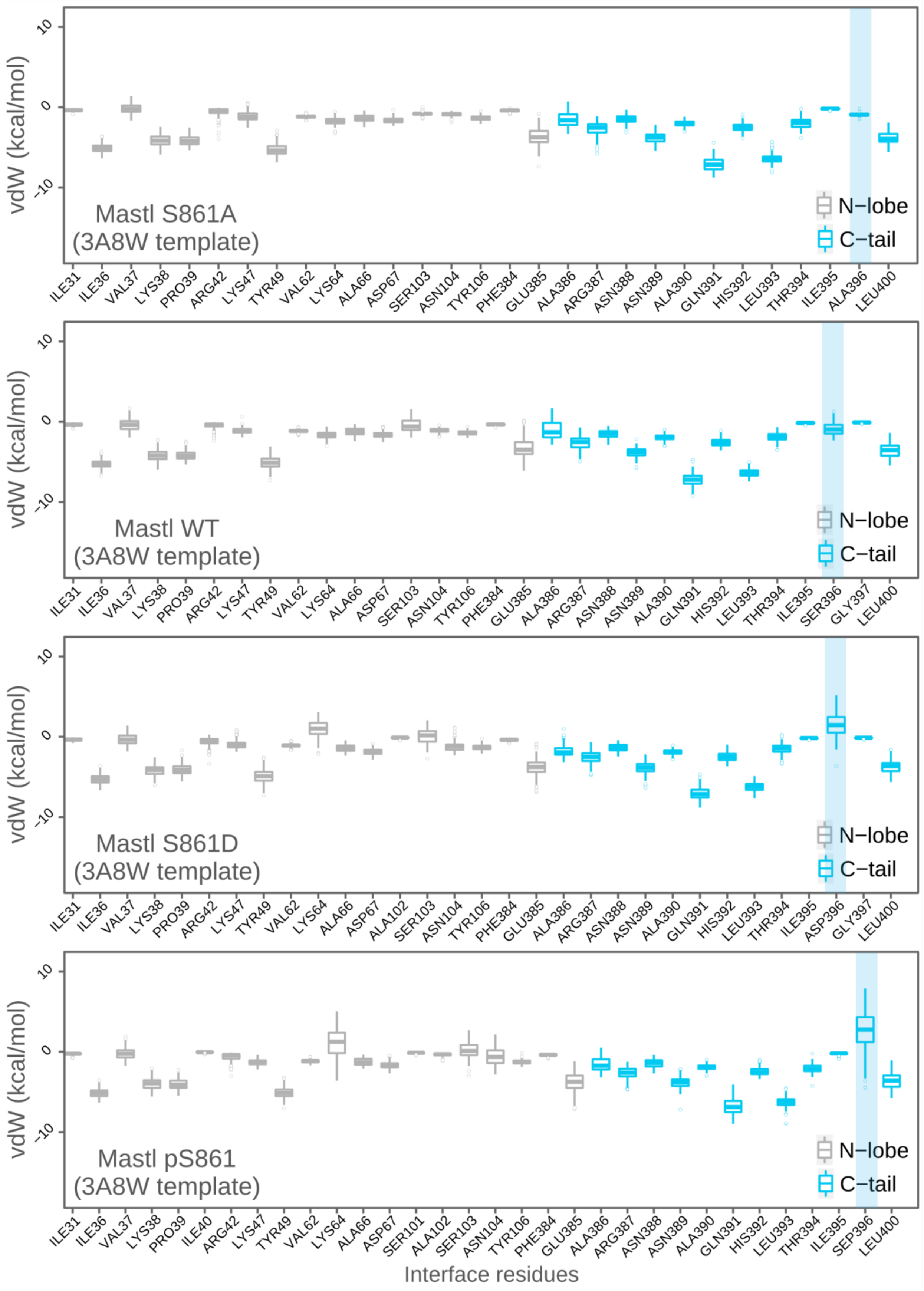
Mastl S861A
(3A8W template)
Mastl WT
(3A8W template)
Mastl S861D
(3A8W template)
Mastl pS861
(3A8W template)
vdW (kcal/mol)
10
0
-10
N−lobe
C−tail
ILE31 ILE36 VAL37 LYS38 PRO39 ARG42 LYS47 TYR49 VAL62 LYS64 ALA66 ASP67 SER103 ASN104 TYR106 PHE384 GLU385 ALA386 ARG387 ASN388 ASN389 ALA390 GLN391 HIS392 LEU393 THR394 ILE395 ALA396 LEU400
ILE31 ILE36 VAL37 LYS38 PRO39 ARG42 LYS47 TYR49 VAL62 LYS64 ALA66 ASP67 SER103 ASN104 TYR106 PHE384 GLU385 ALA386 ARG387 ASN388 ASN389 ALA390 GLN391 HIS392 LEU393 THR394 ILE395 SER396 GLY397 LEU400
ILE31 ILE36 VAL37 LYS38 PRO39 ARG42 LYS47 TYR49 VAL62 LYS64 ALA66 ASP67 ALA102 SER103 ASN104 TYR106 PHE384 GLU385 ALA386 ARG387 ASN388 ASN389 ALA390 GLN391 HIS392 LEU393 THR394 ILE395 ASP396 GLY397 LEU400
ILE31 ILE36 VAL37 LYS38 PRO39 ILE40 ARG42 LYS47 TYR49 VAL62 LYS64 ALA66 ASP67 SER101 ALA102 SER103 ASN104 TYR106 PHE384 GLU385 ALA386 ARG387 ASN388 ASN389 ALA390 GLN391 HIS392 LEU393 THR394 ILE395 SEP396 LEU400
Interface residues

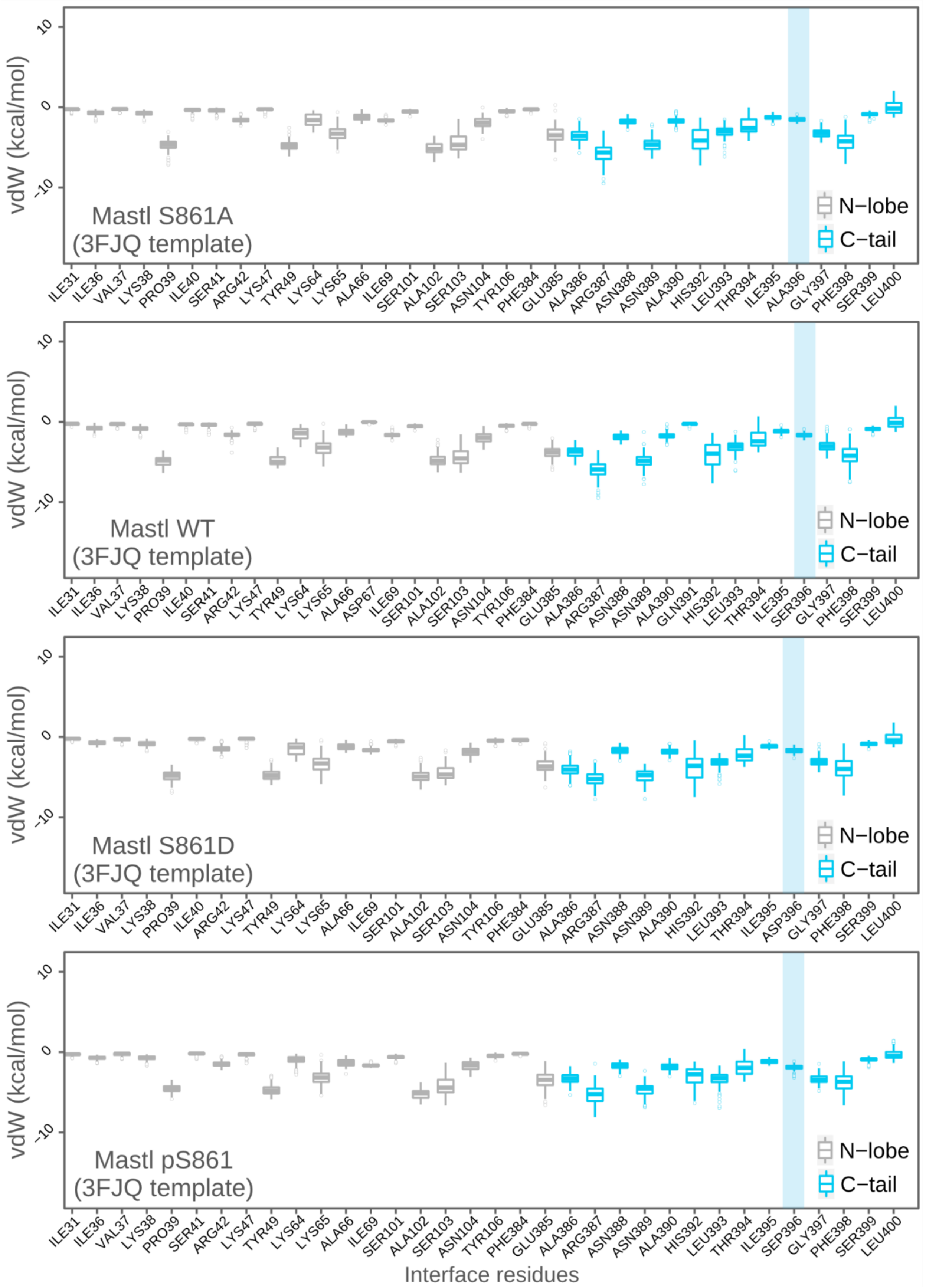
vdW (kcal/mol)
10
0
-10
Mastl S861A
(3FJQ template)
N−lobe
C−tail
ILE31 ILE36 VAL37 LYS38 PRO39 ILE40 SER41 ARG42 LYS47 TYR49 LYS64 LYS65 ALA66 ILE69 SER101 ALA102 SER103 ASN104 TYR106 PHE384 GLU385 ALA386 ARG387 ASN388 ASN389 ALA390 HIS392 LEU393 THR394 ILE395 ALA396 GLY397 PHE398 SER399 LEU400
Mastl WT
(3FJQ template)
ILE31 ILE36 VAL37 LYS38 PRO39 ILE40 SER41 ARG42 LYS47 TYR49 LYS64 LYS65 ALA66 ASP67 ILE69 SER101 ALA102 SER103 ASN104 TYR106 PHE384 GLU385 ALA386 ARG387 ASN388 ASN389 ALA390 GLN391 HIS392 LEU393 THR394 ILE395 SER396 GLY397 PHE398 SER399 LEU400
Mastl S861D
(3FJQ template)
ILE31 ILE36 VAL37 LYS38 PRO39 ILE40 ARG42 LYS47 TYR49 LYS64 LYS65 ALA66 ILE69 SER101 ALA102 SER103 ASN104 TYR106 PHE384 GLU385 ALA386 ARG387 ASN388 ASN389 ALA390 HIS392 LEU393 THR394 ILE395 ASP396 GLY397 PHE398 SER399 LEU400
Mastl pS861
(3FJQ template)
ILE31 ILE36 VAL37 LYS38 PRO39 SER41 ARG42 LYS47 TYR49 LYS64 LYS65 ALA66 ILE69 SER101 ALA102 SER103 ASN104 TYR106 PHE384 GLU385 ALA386 ARG387 ASN388 ASN389 ALA390 HIS392 LEU393 THR394 ILE395 SEP396 GLY397 PHE398 SER399 LEU400
Interface residues

**Figure S7.** The PKC iota crystal structure (3A8W), PKA crystal structure (3FJQ), mouse Mastl 3A8W-based model structure, and mouse Mastl 3FJQ-based model structure were subjected to structural refinement using HADDOCK2.2 web server. For each structure, simulation results of their unphosphorylatable, wild-type, phosphomimetic, and phosphorylated forms are given. Throughout the energy minimization, HADDOCK generated 200 states for each starting structure. For each interface residue, electrostatics score, a hybrid score (vdW + 0.2 x electrostatics score), or van der Waals (vdW) scores of these 200 model structures are given as boxplots. Each box shows the energy distribution of individual interface residues throughout the simulation. The C-tail turn motif and the N-lobe are cyan and gray colored, respectively. The transparent blue rectangle on the right side of each plot indicates the position of the C-tail phosphosite. The residues are numbered according to their positions within the truncated structures. The positions of the phosphosites within the full-length sequences are written in the bottom-left corner of each plot.

**Table S1.** The list of the crystal structures that contain a phosphorylated C-tail. The respective PDB entry Ids, protein names, conformation of the C-tail phosphosites, source of the enzymes, ligand content, and resolution of the structures are given. The structures used for calculations are indicated with gray-filled rows.

| PDB Id | Protein | Phosphoresidue state | Organism | Ligand | Resolution (Å) |
|---|---|---|---|---|---|
| 3A8W | PKCiota | Buried | Hs | ATP | 2.1 |
| 3A8X | PKCiota | Buried | Hs | -none- | 2.00 |
| 1ATP | PKA | Solvent-exposed | Mm | ATP | 2.2 |
| 3FJQ | PKA | Solvent-exposed | Mm | ATP | 1.6 |
| 3X2U | PKA | Solvent-exposed | Mm | ATP | 2.4 |
| 3X2V | PKA | Solvent-exposed | Mm | ATP | 1.77 |
| 3X2W | PKA | Solvent-exposed | Mm | ATP | 1.7 |
| 4DIN | PKA | Solvent-exposed | Mm | ATP | 3.7 |
| 4WB5 | PKA | Solvent-exposed | Hs | ATP | 1.64 |
| 4WB7 | PKA | Solvent-exposed | Hs | ATP | 1.9 |
| 4WB8 | PKA | Solvent-exposed | Hs | ATP | 1.55 |
| 4XW4 | PKA | Solvent-exposed | Mm | ANP | 1.82 |
| 4XW5 | PKA | Solvent-exposed | Mm | ATP | 1.95 |
| 4XW6 | PKA | Solvent-exposed | Mm | ADP | 1.9 |
| 6BYR | PKA | Solvent-exposed | Hs | ATP | 3.66 |
| 6BYS | PKA | Solvent-exposed | Hs | -none- | 4.75 |
| 6NO7 | PKA | Solvent-exposed | Hs | ATP | 3.55 |

**Table S2.** Details of the primers used in this study.

| Purpose | Primer name | Primer sequence (5' to 3') | Size (bp) |
|---|---|---|---|
| PCR genotyping, FOR | KDO227 | CATGCCTTCCTTGAAAGAGGTGGAC | 25 |
| PCR genotyping, REV1 | KDO228 | GTGGGAGGAATTACAAGAGACAAC | 24 |
| PCR genotyping, REV2 | KDO198 | GGCAGGTGGAGGCAAGAGCTCACAGA | 26 |
| Mouse Mastl T192V mutagenesis forward | KDO229 | GGATATTCTCACAGTACCATCAATGTCTAAACC | 33 |
| Mouse Mastl T192V mutagenesis reverse | KDO230 | GGTTTAGACATTGATGGTACTGTGAGAATATCC | 33 |
| Mouse Mastl T205V mutagenesis forward | KDO231 | GATTATTCAAGAGTTCCAGGACAAGTCTTATCTCT | 35 |
| Mouse Mastl T205V mutagenesis reverse | KDO232 | GTCCTGGAACTCTTGAATAATCTTGCTTAGG | 31 |
| Mouse Mastl S211A mutagenesis forward | KDO233 | CAAGTCTTAGCTCTCATCAGCTCTTTGG | 28 |
| Mouse Mastl S211A mutagenesis reverse | KDO234 | GCTGATGAGAGCTAAGACTTGTCCTG | 26 |
| Mouse Mastl T726V mutagenesis forward | KDO235 | TAGGGGTTCCAGATTACCTGGC | 22 |
| Mouse Mastl T726V mutagenesis reverse | KDO236 | CAGGTAATCTGGAACCCCTAGAATTCG | 27 |
| Mouse Mastl S861A mutagenesis reverse | KDO237 | CTGCAGAACCACTGTGCTGGCTACAGACTAAACCCAGCTATGGTCAGATG | 50 |
| Mouse Mastl T192E mutagenesis forward | KDO238 | GGATATTCTCACAGAACCATCAATGTCTAAACC | 33 |
| Mouse Mastl T192E mutagenesis reverse | KDO239 | GGTTTAGACATTGATGGTTCTGTGAGAATATCC | 33 |
| Mouse Mastl T205E mutagenesis forward | KDO240 | GATTATTCAAGAGAACCAGGACAAGTCTTATCTCT | 35 |
| Mouse Mastl T205E mutagenesis reverse | KDO241 | GTCCTGGTTCTCTTGAATAATCTTGCTTAGG | 31 |

| | | | |
|---|---|---|---|
| Mouse Mastl S861D mutagenesis reverse | KDO242 | CTGCAGAACCACTGTGCTGGCTACAGACTA AACCCATCTATGGTCAGATG | 50 |
